# Direct fluorescent labeling of NF186 and Na_V_1.6 in living primary neurons using bioorthogonal click chemistry

**DOI:** 10.1101/2022.03.01.480798

**Authors:** Nevena Stajković, Yuanyuan Liu, Aleksandra Arsić, Ning Meng, Hang Lyu, Nan Zhang, Dirk Grimm, Holger Lerche, Ivana Nikić-Spiegel

**Affiliations:** Werner Reichardt Centre for Integrative Neuroscience, University of Tübingen, Tübingen, Germany; Graduate Training Centre of Neuroscience, International Max Planck Research School, University of Tübingen, Tübingen, Germany; Department of Neurology and Epileptology, Hertie Institute for Clinical Brain Research, University of Tübingen, Tübingen, Germany; Virus-Host Interaction Group, Department of Infectious Diseases/Virology, Medical Faculty, University of Heidelberg, Cluster of Excellence CellNetworks, BioQuant, Heidelberg, Germany; German Center for Infection Research and German Center for Cardiovascular Research, partner site Heidelberg, Heidelberg, Germany

## Abstract

The axon initial segment (AIS) is a highly specialized neuronal compartment that regulates the generation of action potentials and maintenance of neuronal polarity. Despite its importance, live imaging of the AIS is challenging due to the limited number of suitable labeling methods. To overcome this limitation, we established a novel approach for live labeling of the AIS using unnatural amino acids (UAAs) and bioorthogonal click chemistry. The small size of the UAAs and the possibility of introducing them virtually anywhere into the target proteins make this method particularly suitable for live labeling and imaging of complex and spatially restricted proteins. With this approach, we labeled two large AIS components, the 186 kDa isoform of neurofascin (NF186) and the 260 kDa voltage-gated sodium channel (Na_V_1.6), and performed widefield and confocal microscopy in fixed and living neurons. Moreover, we demonstrated the applicability of this method by studying the localization of two epilepsy-causing Na_V_1.6 variants with a loss-of-function effect. Finally, to further improve the efficiency of the UAA incorporation, we developed adeno-associated viral (AAV) vectors for click labeling in primary neurons. The use of AAV vectors will facilitate the transfer of UAA-based click labeling technology to more complex biological systems, such as organotypic slice cultures, organoids, and animal models.

## Introduction

The axon initial segment (AIS) is a highly specialized neuronal compartment responsible for the generation of action potentials (Leterrier, 2018). This unique role of the AIS is mediated by high-density accumulations of voltage-gated ion channels in it (Kole et al., 2008). Particularly important among these channels is Na_V_1.6, the most abundant voltage-gated sodium channel isoform in the adult mammalian brain (Sole and Tamkun, 2020). Na_V_1.6 clusters at the distal AIS (Hu et al., 2009) through interactions with the membrane domain of ankyrin G (ankG) (Leterrier, 2018). AnkG acts as an adaptor that anchors Na_V_1.6 and other important AIS components, such as the 186 kDa neurofascin isoform (NF186), to the underlying cytoskeleton (Leterrier, 2018). As revealed by super-resolution microscopy, AIS constituents are evenly spaced along the AIS with a periodicity of ^~^190 nm (Leterrier et al., 2015, Xu et al., 2013).

Correct function and subcellular targeting of Na_V_1.6 and NF186 are crucial for proper neuronal activity. Consequently, genetic variations and secondary alterations of Na_V_1.6 have been implicated in various neurological diseases, such as epilepsy, autism, intellectual disability, movement disorders, and multiple sclerosis (Craner et al., 2004, Johannesen et al., 2021, Meisler et al., 2021), while auto-antibodies against NF186 have been found in multiple sclerosis and chronic inflammatory demyelinating polyradiculoneuropathy (Kira et al., 2019).

To study the trafficking and dynamics of Na_V_1.6 and NF186, several live-cell labeling approaches have been developed (Akin et al., 2015, Akin et al., 2016, Dzhashiashvili et al., 2007, Gasser et al., 2012, Ghosh et al., 2020, Liu et al., 2021, Sole et al., 2019, Susuki et al., 2013, Zhang et al., 1998). The most widely used approach relies on generating genetic fusions with fluorescent proteins (FPs; **Fig. 1a**) (Akin et al., 2015, Dzhashiashvili et al., 2007, Gasser et al., 2012, Ghosh et al., 2020, Zhang et al., 1998). The main advantages of FP fusions are their high specificity and compatibility with live-cell imaging. However, most fusions are made by placing the FP at either the N or C terminus of the target protein. When it comes to the AIS components, these domains frequently participate in channel inactivation, targeting, and localization, or include binding regions for different regulatory proteins. Some of these interactions may be impaired by relatively large FP tags (^~^30 kDa). For example, it has been reported that the addition of a green fluorescent protein (GFP) to the C terminus of NF186 results in its mislocalization (Dumitrescu et al., 2016, Dzhashiashvili et al., 2007). In addition to FP fusions, additional methods have been established for live labeling of Na_V_1.6 and other voltage-gated sodium channel isoforms. One of the approaches for specific labeling of Na_V_ channels is based on the incorporation of a 17-amino-acid-long biotinylated domain (BAD) into their extracellular domains. The BAD domain, when biotinylated by bacterial biotin ligase, can be labeled with non-permeable streptavidin conjugated dyes. This method has been successfully used to label Na_V_1.6 (Akin et al., 2015), Na_V_1.9 (Akin et al., 2019), and Na_V_1.7 (Sizova et al., 2020) in primary neurons or mammalian cell lines. However, the disadvantages of this approach are the bulkiness and the large size of streptavidin (^~^60 kDa). Recently, a self-labeling HaloTag fused to the C termini of endogenous Na_V_1.6 and Na_V_1.2 has been utilized for live-cell microscopy studies (Liu et al., 2021). This CRISPRC/Cas9 genome editing-based labeling approach offers a great opportunity to study trafficking and localization of endogenous sodium channels, but due to the large size of the HaloTag (^~^33 kDa), this approach has disadvantages that are similar to those of FPs.

**Fig. 1.**
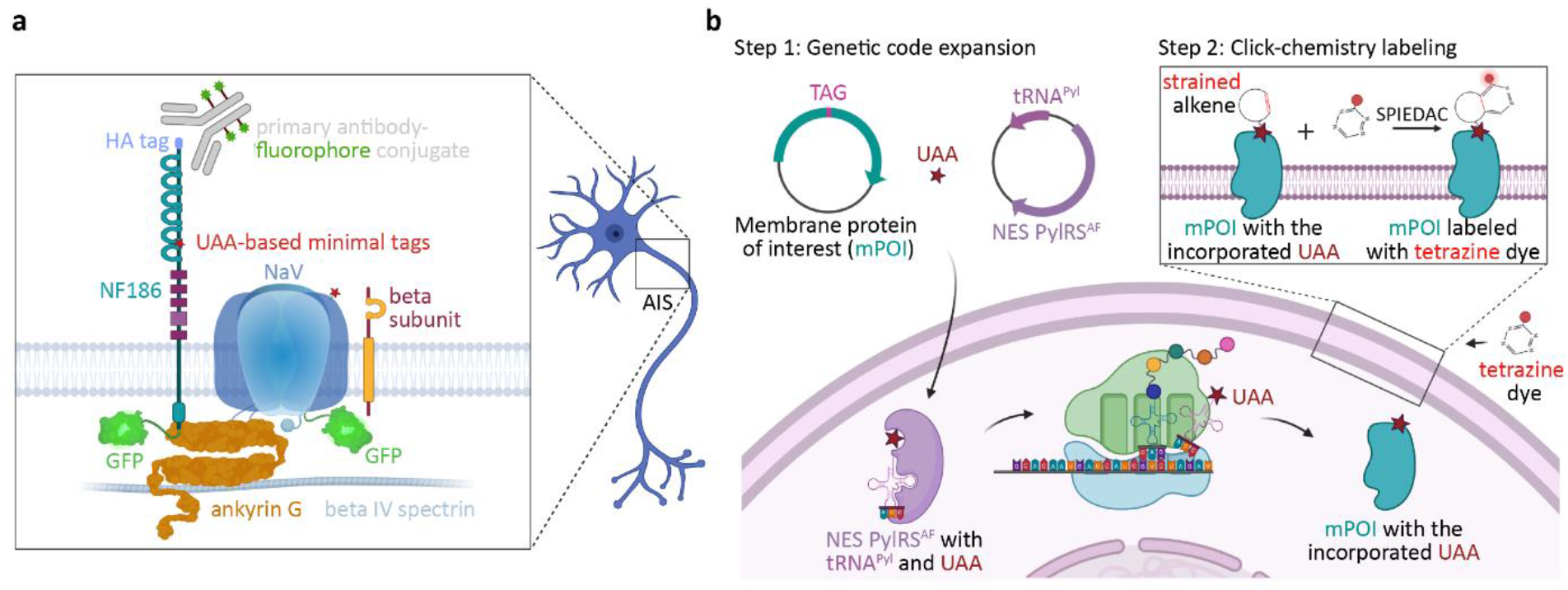
Schematic overview of the methods for live-cell labeling of the axon initial segment (AIS), including unnatural amino acid (UAA)-based minimal tags. **a**: Fluorescent protein (FP)-based tags fused to the C terminus are widely used for live labeling of membrane proteins, such as NF186 (Dzhashiashvili et al., 2007) and voltage-gated sodium channels (Akin et al., 2015; Gasser et al., 2012) or submembrane proteins such as ankyrin G (Jenkins et al., 2015). Alternatively, antibodies conjugated to fluorescent dyes can be utilized for live labeling of extracellular domains of membrane proteins, such as NF186 (Evans et al., 2015; Schafer et al., 2009; Torii et al., 2020). As we showed in this study, UAAs and click chemistry can also be used as minimal tags for live-cell labeling of AIS components. **b**: Schematic representation of genetic code expansion (amber codon suppression) and click labeling of transmembrane proteins. In Step 1, mammalian cells were transfected with plasmids bearing genes that encode a membrane protein of interest [mPOI; that had been mutated to contain an in-frame amber (TAG) stop codon], the Y306A/Y384F (AF) double-mutant of the orthogonal pyrrolysyl (Pyl) tRNA synthetase fused to a nuclear export signal (NES PylRS^AF^), and its cognate amber codon suppressor tRNA^Pyl^ derived from *Methanosarcina mazei*. During the protein translation, the NES PylRS^AF^/tRNA^Pyl^ pair enabled site-specific incorporation of the UAA, such as trans-c*yclooct-2-en*-L-lysine (TCO*A-Lys), into the mPOI in response to the TAG stop codon. Subsequently (Step 2), TCO*A-Lys was coupled to a cell-impermeable fluorescent tetrazine dye in a strain-promoted inverse-electron demand Diels-Alder cycloaddition (SPIEDAC) click reaction. The schemes were made in BioRender.com.

Alternatively to genetic fusions, immunostaining with fluorescent antibodies can be used for AIS labeling in living neurons. For this purpose, antibodies that recognize either extracellular domains of endogenous AIS proteins or short tags attached to the extracellular domains of recombinant/endogenous AIS components have been utilized (**Fig. 1a**) (Dumitrescu et al., 2016, Dzhashiashvili et al., 2007, Evans et al., 2015, Freal et al., 2019, Hedstrom et al., 2008, Liu et al., 2021, Schafer et al., 2009, Torii et al., 2020). However, antibodies, due to their multivalence, can induce crosslinking, which makes them unsuitable to study the dynamics of the AIS, such as plasticity (Dumitrescu et al., 2016). Furthermore, when it comes to the labeling of closely-related proteins such as different Na_V_ isoforms, antibodies do not always provide enough specificity. Keeping these limitations in mind, it would be beneficial to develop additional approaches for direct and minimally invasive live-cell labeling of AIS components.

We and other researchers previously developed unnatural amino acid (UAA)-based minimal tags for live-cell protein labeling in mammalian cells (Lang et al., 2012b, Lang et al., 2012a, Nikic et al., 2014, Plass et al., 2012, Uttamapinant et al., 2015). These tags are based on site-specific labeling of a single UAA with a fluorescent dye. With the help of genetic code expansion (Chin, 2017, Nikic-Spiegel, 2020, Wang, 2017), UAAs carrying strained alkenes are co-translationally incorporated into the protein of interest in response to an in-frame amber stop codon (**Fig. 1b**). Subsequently, fluorescent dyes are directly attached to the UAAs with click chemistry reactions. One such reaction is the bioorthogonal catalyst-free strain-promoted inverse-electron demand Diels-Alder cycloaddition (SPIEDAC) between UAAs that contain strained alkenes and tetrazine derivatives of fluorescent dyes. Due to its high reaction rates and biorthogonality, SPIEDAC is particularly useful for live-cell labeling. In recent years, this type of labeling has emerged as one of the most powerful approaches for the minimally invasive labeling of both extracellular and intracellular proteins in standard cell lines. We have recently established this type of labeling in primary neurons by labeling the neuronal cytoskeleton (Arsić et al., 2022). In parallel, this approach has been used to label small (34–45 kDa) transmembrane AMPA receptor regulatory proteins (Bessa-Neto et al., 2021).

In this study, we utilized SPIEDAC to label two large transmembrane AIS proteins: NF186 (^~^186 kDa) and Na_V_1.6 (^~^260 kDa). Direct fluorescent labeling of the UAA with small dyes allowed us to perform fixed and live-cell confocal microscopy and to study localization of WT and pathogenic Na_V_1.6 variants. We also developed adeno-associated viral (AAV) vectors that mediated high-efficiency delivery of the genetic code expansion components to primary neurons.

## Results

### Genetic code expansion and click labeling of NF186 in the ND7/23 cell line

To establish live-cell labeling of the AIS using click chemistry, we first focused on the labeling of one of its smaller (186 kDa) components, NF186 (**Fig. 1a**). Before introducing amber codon (TAG) labeling sites via site-directed mutagenesis, we modified the previously described plasmid (Zhang et al., 1998) by moving the hemagglutinin (HA) tag from the N terminus to the C terminus. Since this plasmid encodes the wild-type (WT) rat NF186 driven by a cytomegalovirus (CMV) promoter, we will refer to it hereafter as a “CMV-NF186^WT^-HA” construct. The C-terminal HA-tag allowed us to detect the full-length NF186 protein by immunostaining it with an anti-HA antibody. To find the optimal TAG mutant in terms of expression and click labeling efficiency, we tested different TAG positions in NF186. Testing different positions for click labeling of a target protein is an important step, considering that the amber codon suppression efficiency depends on the surrounding sequence (Bartoschek et al., 2021), while the labeling efficiency depends on the accessibility of a UAA to a tetrazine dye. SWISS-MODEL (Bienert et al., 2017), based on the crystal structure of a titin fragment (PDB 3B43) (von Castelmur et al., 2008) was used to select six potential positions for the UAA incorporation in the extracellular domain of NF186 (**Fig. 2a**). Then, we introduced the corresponding amber (TAG) codons into the rat *Nfasc* gene by site-specifically mutating the following lysine (K) residues: NF186^K519TAG^, NF186^K534TAG^, NF186^K571TAG^, NF186^K604TAG^, NF186^K680TAG^, and NF186^K809TAG^.

**Fig. 2.**
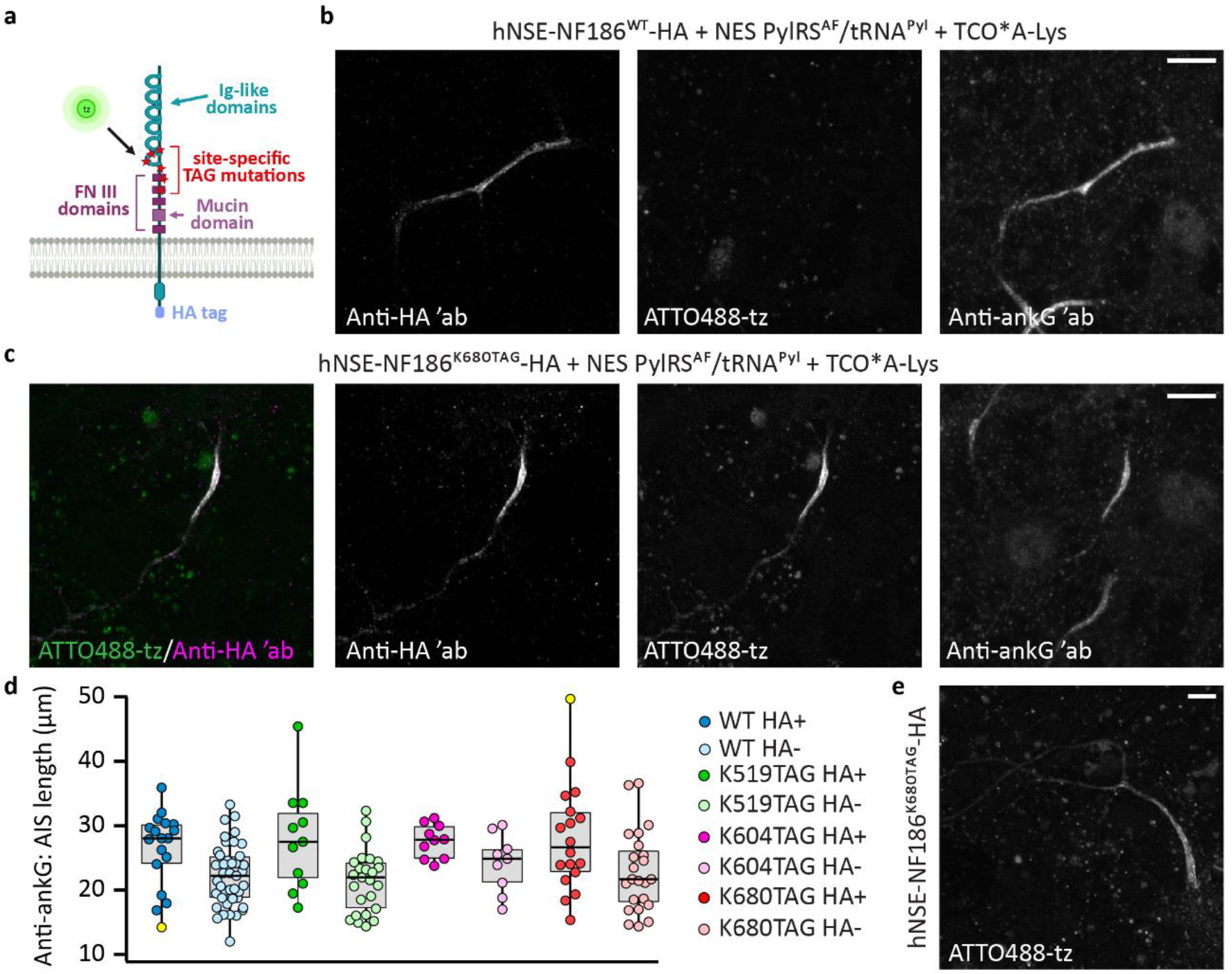
Genetic code expansion and click labeling of NF186 in living primary neurons. **a:** Schematic representation of the NF186 structure with clickable labeling sites (red) and the C-terminal hemagglutinin (HA) tag (purple). **b**–**c**: Representative confocal images of primary rat cortical neurons at day *in vitro* (DIV) 11 expressing NES PylRS^AF^/tRNA^Pyl^ and NF186^WT^-HA or NF186^K680TAG^-HA under the human neuron-specific enolase (hNSE) promoter, in the presence of the unnatural amino acid TCO*A-Lys. Prior to imaging, living neurons were click-labeled with ATTO488-tetrazine (tz), fixed, and immunostained with anti-HA and anti-ankyrin G (ankG) primary antibodies, followed by Alexa Fluor (AF) 555- and AF633-conjugated secondary antibodies. **d**: Distribution of the AIS lengths measured in confocal images of the anti-ankG immunostained rat neurons expressing NF186^WT^-HA, NF186K519TAG-HA, NF186^K604TAG^-HA, or NF186^K680TAG^-HA (HA+) and the corresponding neighboring untransfected (HA-) neurons. The box plots indicate the median (the black lines inside the box), the 25^th^ and 75^th^ percentiles (the box boundaries), the single data points (the dots), and the outliers (the yellow dots). The whiskers lengths are defined by the minimum and maximum data points. The non-parametric Kruskal-Wallis test with Dunn-Bonferroni *posthoc* did not show any significant differences between the HA+ and HA-AIS lengths for WT and the three clickable mutants (p_WT_ = 0.206, p_K519TAG_ = 0.265 p_K604TAG_ = 1.000, and p_K680TAG_ = 0.605; number (n) of analyzed cells: n_WT HA+_ =18, n_WT HA-_ = 41, n_K519TAG HA+_ = 11, n_K519TAG HA-_ = 25, n_K604TAG HA+_ = 10, n_K604TAG HA-_ = 9, n_K680TAG HA+_ = 18, and n_K680TAG HA-_ = 23). The details of the statistical analysis are shown in **Supplementary Table 1**. **e**: Representative confocal image of a living neuron at DIV 11 expressing NF186^K680TAG-^HA labeled with ATTO488-tz. The Z-stack images are shown as maximum intensity projections in all panels. Scale bars: 10 μm **(b**, **c**, **e)**. The scheme in panel a was made in BioRender.com.

We have recently shown that the rodent neuroblastoma ND7/23 cell line is a suitable host for genetic code expansion and click labeling of neuronal proteins (Arsić et al., 2022). To identify the optimal TAG position for NF186 click labeling, we co-transfected ND7/23 cells with CMV-NF186^TAG^-HA constructs and the previously described codon-optimized plasmid for clickable UAA incorporation. The latter encodes the Y306A/Y384F (AF) double mutant of the *Methanosarcina mazei-derived* orthogonal pyrrolysyl (Pyl) tRNA synthetase fused to a nuclear export signal (NES PylRS^AF^) and its cognate amber codon suppressor tRNA^Pyl^ (Arsić et al., 2022). The immunostaining with an anti-HA antibody and confocal imaging confirmed that all the mutants were expressed in the presence of the trans-cyclooct-2-en-L-lysine (TCO*A-Lys) UAA, whereas no expression was detected in the absence of TCO*A-Lys (**Supplementary Fig. 1 and 2**). Furthermore, live-cell click labeling with an ATTO488-tetrazine derivative (ATTO488-tz) showed that all six clickable TAG mutants can be labeled, albeit with different efficiencies (**Supplementary Fig. 1**). Expectedly, in the absence of UAA, there was no click labeling (**Supplementary Fig. 2**).

### Genetic code expansion and click labeling of NF186 in living primary neurons

Having established click labeling of NF186^TAG^-HA in the ND7/23 cell line, we aimed to implement labeling of NF186 in primary rat cortical neurons. Typically, we would pick the best expressing TAG mutant with the highest labeling efficiency from the ND7/23 screen, since this construct has the highest chance of being successfully expressed and labeled in primary neurons. However, it is well established that overexpression of certain AIS components such as ankG and NF186 can lead to their mislocalization or can result in an abnormally elongated AIS (Dumitrescu et al., 2016, Galiano et al., 2012, Hamdan et al., 2020, Jenkins et al., 2015). Considering that the NF186^TAG^-HA mutants from our ND7/23 screen had a lower expression than the NF186^WT^-HA, we anticipated that this would overcome mislocalization problem. For confirmation, we tested all six NF186^TAG^-HA mutants in the primary neurons (**Supplementary Fig. 3**). Four days after the transfection, on day *in vitro* (DIV) 11, we performed immunostaining with an anti-HA antibody to estimate the expression level and localization of the recombinant NF186. As expected, the NF186^WT^-HA expressed from a strong CMV promoter was mostly mislocalized and expressed in the soma and other neuronal processes (**Supplementary Fig. 3a**). NF186^TAG^-HA mutants, despite their lower expression levels, also frequently showed ectopic localization along distal axons (**Supplementary Fig. 3b**).

To improve the localization of the NF186-HA constructs, we replaced the CMV promoter with the weaker human neuron-specific enolase promoter (hNSE) in the NF186^WT^-HA and NF186^TAG^-HA constructs (hereafter referred to as “hNSE-NF186^WT^-HA” and “hNSE-NF186^TAG^-HA”). This promoter was previously used to lower the expression level of NF186 in hippocampal neurons (Hamdan et al., 2020). Four days after the transfections, we performed click labeling with ATTO488-tz, followed by immunostaining with anti-HA and anti-ankG antibodies (**Fig. 2b-c**, **Supplementary Fig. 4**). Confocal microscopy revealed that the hNSE promoter lowered the WT and clickable NF186-HA expression levels and consequently improved the localization of these proteins (**Fig. 2b-c and Supplementary Fig. 4**). Furthermore, confocal microscopy confirmed that only the hNSE-NF186^TAG^-HA constructs and not hNSE-NF186^WT^-HA could be labeled with click chemistry. Finally, these experiments revealed that not all the NF186^TAG^ mutants could be equally well labeled with ATTO488-tz in primary neurons (**Fig. 2c and Supplementary Fig. 4**). For example, the number of transfected neurons with NF186^K534TAG^-HA and NF186^K571TAG^-HA constructs was lower, and click labeling was either weak or completely absent (**Supplementary Fig. 4**). Therefore, we excluded these two mutants from our further analysis. Although NF186^K809TAG^-HA (**Supplementary Fig. 4**) showed bright click labeling, we excluded it from the analysis due to its frequent ectopic expression along the distal axon.

To identify the most suitable TAG position for NF186 click labeling among the three remaining mutants (NF186^K519TAG^-HA, NF186^K604TAG^-HA, and NF186^K680TAG^-HA), we assessed whether the AIS structure was affected by overexpression of the hNSE-NF186^TAG^-HA constructs. To this end, we used the previously custom-written MATLAB script for quantitative measurements of the AIS length (Grubb, 2021, Grubb and Burrone, 2010) to compare the AIS length of the NF186^WT/TAG^-HA transfected (HA positive) and neighboring untransfected (HA negative) neurons (**Fig. 2d**). Anti-HA immunostaining allowed us to identify transfected neurons, while we used immunostaining with anti-ankG as a transfection-independent marker to identify the AIS in both transfected and surrounding untransfected neurons. Quantitative analysis of the AIS length showed no significant differences between the AIS length of the neurons that expressed recombinant NF186^WT^-HA or clickable NF186^TAG^-HA and the AIS length of the untransfected neurons (**Fig. 2d**).

Considering that neither of the three analyzed NF186^TAG^-HA amber mutants affected the AIS length, we used an additional parameter to determine the most suitable mutant for click labeling of NF186 in primary neurons. Among other factors, the UAA incorporation efficiency depends on the sequence surrounding the TAG amber codon (Bartoschek et al., 2021). We used a recently developed tool (Bartoschek et al., 2021) to predict the UAA incorporation efficiency rate in the three remaining amber mutants. The iPASS (identification of <u>p</u>ermissive <u>a</u>mber <u>s</u>ites for <u>s</u>uppression) score was higher for NF186^K680TAG^-HA (1.83) than for NF186^K519TAG^-HA (1.61) and NF186^K604TAG^-HA (1.63). Considering the high iPASS score and the reproducibly high labeling intensity of NF186^K680TAG^-HA, we decided to use it in future experiments. This allowed us to perform fixed-cell (**Fig. 2c**) and live-cell confocal imaging of click-labeled NF186 in primary neurons (**Fig. 2e**). Altogether, these results demonstrate successful application of genetic code expansion and click chemistry for the labeling and imaging of the AIS in living primary neurons.

### Click labeling of Na_V_1.6 channels in living primary neurons

Once we had established click labeling of NF186 in living primary neurons, we focused on labeling of the alpha subunit of the voltage-gated sodium channel isoform Na_V_1.6 (encoded by the *SCN8A* gene). Considering the highly complex structure of Na_V_1.6 that involves folding a single polypeptide chain of ^~^2000 amino acids (**Fig. 3a**) into four homologous domains (I to IV), each of which contains six transmembrane segments (S1 to S6), we had to address additional challenges while selecting potential positions for UAA incorporation. First of all, there are many disease-related mutations in the *SCN8A* gene (Johannesen et al., 2021, Meisler et al., 2021), and there are many conserved regions of Na_V_1.6 that are crucial for the function of the channel (e.g., the S4 segments of domains I–IV and pore-forming loops; **Fig. 3a**) (Catterall et al., 2005). These positions/domains had to be avoided when selecting TAG positions to ensure that the function of the Na_V_1.6 would not be affected. Additionally, the crystal structure of Na_V_1.6 is not available, and the alpha subunit of the channel is heavily glycosylated, which made it harder to select TAG positions that would be accessible for the tetrazine dyes. Based on the available literature (Akin et al., 2015, Meisler et al., 2021, O’Brien and Meisler, 2013), we selected two positions (K1425 and K1546) in the extracellular loops of Na_V_1.6. These positions are located in less conserved regions of the channel, do not participate in the pore formation, opening of the channel, or the regulation of its function; and are not related to any known diseases. We introduced respective TAG mutations into the corresponding sites of a plasmid encoding WT mouse Na_V_1.6 (**Fig. 3a**). We also added an HA tag to the C terminus of Na_V_1.6, which allowed us to detect full-length proteins and to assess Na_V_1.6 localization in transfected cells and primary neurons.

**Fig. 3.**
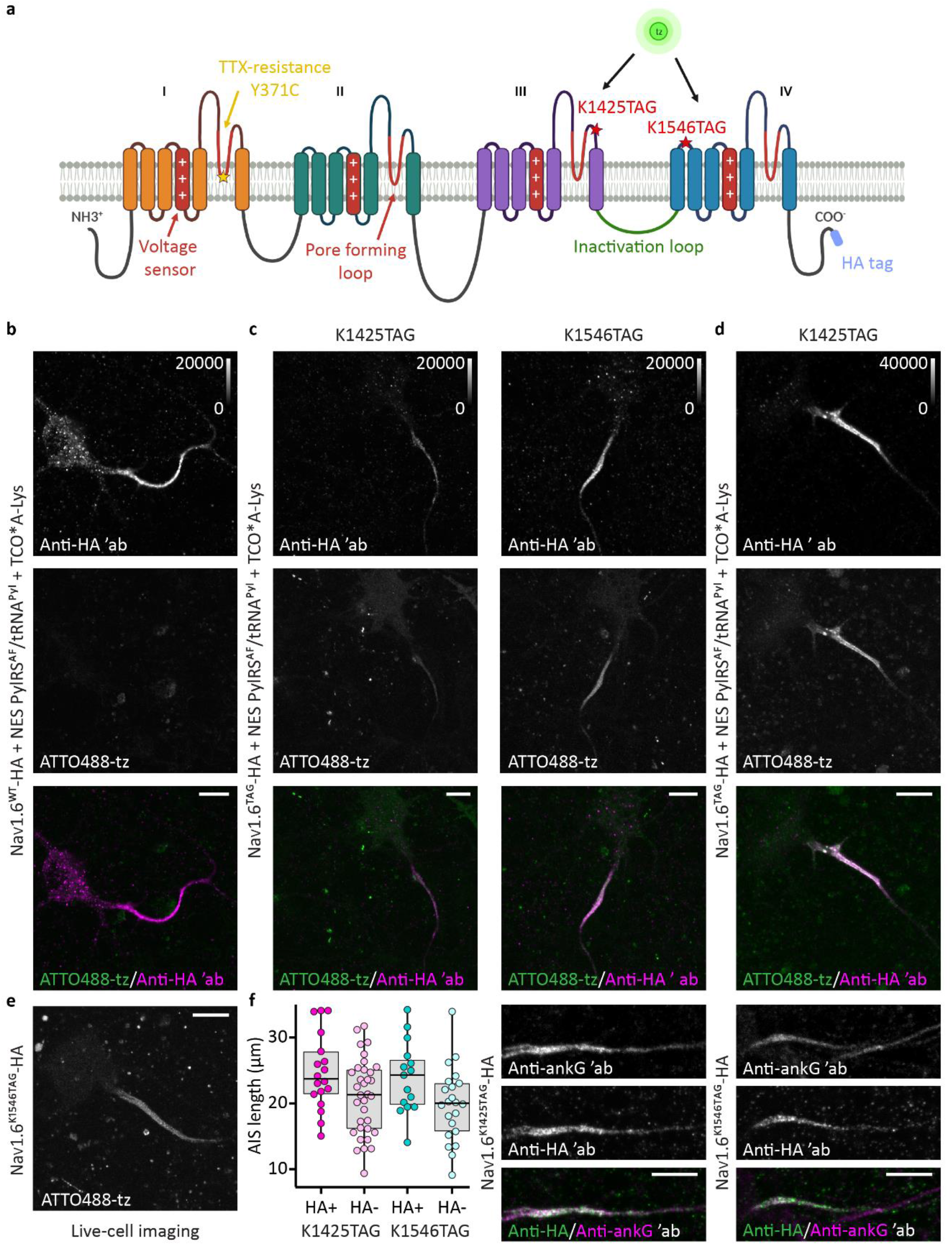
Genetic code expansion and click labeling of Na_V_1.6 in living primary neurons. **a**: Schematic representation of the Na_V_1.6 alpha subunit with the clickable labeling sites (K1425TAG and K1546TAG; red stars) and the C-terminal hemagglutinin (HA) tag (purple). In addition, the scheme depicts the Y371C mutation (yellow star) that rendered Na_V_1.6 tetrodotoxin (TTX)-resistant. **b**–**d**: Representative confocal images of primary rat cortical neurons at DIV 12 expressing NES PylRS^AF^/tRNA^Pyl^ and Na_V_1.6^WT^-HA, Na_V_1.6^K1425TAG^-HA, or Na_V_1.6^K1546TAG^-HA in the presence of the UAA TCO*A-Lys. The neurons were transfected with either Lipofectamine 2000 (b,c,e) or Lipofectamine 3000 (d) reagent. Prior to imaging, living neurons were click-labeled with ATTO488-tetrazine (tz), fixed, and immunostained with an anti-HA primary antibody, followed by an Alexa Fluor 555-conjugated secondary antibody. **e**: Representative confocal image of a living rat neuron at DIV 12 expressing Na_V_1.6^K1546TAG^-HA labeled with ATTO488-tz. **f**: Distribution of the AIS lengths measured in confocal images of anti-ankG immunostained rat neurons expressing Na_V_1.6^K1425TAG^-HA or Na_V_1.6^K1546TAG^-HA (HA+) and the corresponding neighboring untransfected (HA-) neurons. The box plots indicate the median (the black lines inside the box), the 25^th^ and 75^th^ percentiles (the box boundaries), and the single data points (the dots). The whiskers lengths are defined by the minimum and maximum data points. One-way ANOVA with Tukey *posthoc* did not show significant differences between the HA+ and HA-AIS lengths for both clickable mutants (p_K1425TAG_ = 0.124, p_K1546TAG_ = 0.136; number (n) of analyzed cells: n_K1425TAG HA+_ = 18, n_K1425TAG HA-_ = 34, n_K1546TAG HA+_ = 15, and n_K1546TAG HA-_ = 22). The details of the statistical analysis are shown in **Supplementary Table 2**. Representative images of rat neurons immunostained with anti-ankG and anti-HA antibodies used for the quantitative analysis are shown in panel f. All images, except for that of Na_V_1.6^WT^-HA (shown in panel b), were taken as Z-stacks and are shown as maximum intensity projections. For the comparison of the Lipofectamine 2000 transfection reagent and Lipofctamine 3000, the brightness and contrast of the panels showing the HA channels (**b**-**d**) were linearly adjusted as indicated by the look-up-table (LUT) intensity scale bar. LUT intensity scale bars show the minimum and maximum grey values. Scale bars: 10 μm (**b**–**f**). The scheme in panel a was made in BioRender.com.

Similar to the experiments with NF186, prior to the click labeling of Na_V_1.6 in primary neurons, we first wanted to establish and optimize click labeling conditions of Na_V_1.6^TAG^-HA in neuronal cell lines (**Supplementary fig. 5a**). We initially probed ND7/23 cells that had been widely used for electrophysiological recordings of Na^+^ currents derived from the recombinant tetrodotoxin (TTX)-resistant variants of Na_V_1.6 (Meisler et al., 2021, Sharkey et al., 2009). However, our microscopy experiments revealed that the expression level of Na_V_1.6^WT^-HA (**Supplementary Fig. 5a**) and both Na_V_1.6^TAG^-HA mutants (data not shown) on the membrane of the ND7/23 cells was low. Since the transfection efficiency was low and most of the channels remained inside the cytoplasm, extracellular labeling with the cell-impermeable ATTO488-tz signal was not successful (data not shown). To complement these results functionally, we performed whole-cell patch clamping. We first rendered the WT and clickable Na_V_1.6-HA constructs TTX-resistant by introducing the previously described Y371C mutation (**Fig. 3a**) (Leffler et al., 2005, Liu et al., 2019). Most of the measured ND7/23 cells exhibited peak Na^+^ current amplitudes of less than 0.5 nA in the presence of TTX (data not shown), confirming the microscopy results.

To obtain higher levels of expression, we then tested click labeling conditions in the murine neuroblastoma N1E-115-1 cells (**Supplementary Fig. 5b-d**). Immunostaining with anti-HA antibody revealed that the expression of Na_V_1.6^WT^-HA on the membrane of the N1E-115-1 cells was higher than on the ND7/23 cells (**Supplementary Fig. 5a-c**). However, click labeling of both Na_V_1.6^K1425TAG^-HA and Na_V_1.6^K1546TAG^-HA with ATTO488-tz was not successful (**Supplementary fig. 5d**) indicating insufficient expression of the clickable constructs.

We then reasoned that neuroblastoma cell lines may not be the ideal hosts for click labeling and microscopy studies of Na_V_1.6^TAG^ and that a more native environment is required. Thus, we attempted to express and click label Na_V_1.6^TAG^-HA in primary neurons (**Fig. 3b-d and Supplementary Fig. 6**). Four days after the transfection, on DIV12, we click labeled living neurons with ATTO488-tz dye and performed immunostaining with an anti-HA antibody. As expected, the WT protein was expressed in the presence and in the absence of the UAA, and click labeling was not detected (**Fig. 3b, Supplementary Fig. 6a**).

Furthermore, we observed that both Na_V_1.6^TAG^-HA mutants were successfully expressed and click-labeled in the AIS of the neurons (**Fig. 3c**). In the absence of the UAA, click labeling of amber mutants was not detected (**Supplementary Fig. 6b**). As shown in the representative images in **Fig. 3c**, we observed that the expression level and click labeling intensity of Na_V_1.6^K1425TAG^-HA were lower than those of Na_V_1.6^K1546TAG^-HA. By changing the transfection reagent from Lipofectamine 2000 to Lipofectamine 3000, we improved the expression level and consequently, the labeling intensity of Na_V_1.6^K1425TAG^-HA (**Fig. 3d**). However, even though the expression level increased, the number of transfected neurons remained low. In this regard, we noticed that the number of neurons transfected with Lipofectamine 3000 was lower than that of the neurons transfected with Lipofectamine 2000 (5 vs. 15 neurons per well of an eight-well Lab-Tek II chambered cover glass). The high labeling efficiency of Na_V_1.6^K1546TAG^-HA facilitated the confocal imaging of the living click-labeled neurons (**Fig. 3e**). In conclusion, we successfully applied a combination of genetic code expansion and click chemistry for fluorescent labeling and imaging of the voltage-gated sodium channel isoform Na_V_1.6 in living primary neurons.

### Functional characterization of clickable Na_V_1.6^TAG^-HA amber mutants and effect of their expression on the AIS organization

We next investigated if the AIS structure would be affected by overexpression of the clickable Na_V_1.6 variants. To this end, we compared the AIS length of the Na_v_1.6^TAG^-HA transfected neurons with that of surrounding untransfected neurons as described above for NF186 (Grubb, 2021, Grubb and Burrone, 2010). As shown in **Fig. 3f**, the AIS length of neurons expressing Na_V_1.6^TAG^-HA (HA positive) mutants did not significantly differ from that of the untransfected (HA negative) neurons.

Furthermore, we wanted to investigate if incorporation of TCO*A-Lys into Na_V_1.6 and click labeling had an impact on biophysical properties of the channels or on the nanoscale organization of the AIS. To assess whether functional properties of the UAA-tagged channels are affected, we performed whole-cell patch clamp recordings of Na^+^ currents in N1E-115-1 cells (**Fig. 4a-d and Supplementary Fig. 7**). Those were co-transfected with TTX-resistant Na_V_1.6^WT, Y371C^-HA or clickable Na_V_1.6^K1546TAG^, ^Y371C^-HA plasmids, NES PylRS^AF^/tRNA^Pyl^, and a multigene plasmid that contained mouse β1- and β2-subunits and GFP to ensure full functionality of Na_V_1.6 channels and to identify transfected cells, respectively. The K1546TAG mutation caused a small but significant depolarizing shift (2.8 mV) of the fast inactivation curve, slightly slowed the time course of fast inactivation and accelerated its recovery (**Fig. 4a and c; Supplementary Fig. 7a-e**; **Supplementary Table 03**). For the Na_V_1.6^K1425TAG, Y371C^-HA variant (**Fig. 4b and d; Supplementary Fig. 7f-j**), the number of transfected cells and the expression level were lower than those of Na_V_1.6^K1546TAG, Y371C^-HA, corresponding to the reduced Na^+^ currents. To increase the transfection efficiency, we used the commercial PiggyBAC transposase system to stably incorporate the mouse β1- and β2-subunits into the genome of the N1E-115-1 cells reducing the number of plasmids needed for transfections of the N1E-115-1 cells. To identify the transfected cells, we generated a plasmid containing the genes encoding Na_V_1.6^Y371C^ and enhanced GFP (eGFP), separated by a self-cleaving P2A sequence (Na_V_1.6^Y371C^-P2A-eGFP). Under these conditions, we acquired a sufficient number of transfected cells and recorded larger Na^+^ currents. The K1425TAG clickable variant reduced the peak current density compared to the WT channel, whereas changes of other gating parameters were not observed (**Fig. 4b and d; Supplementary Fig. 07f-j, Supplementary Table 3**). The reduction of the peak current density corresponds well to the reduced expression level of Na_V_1.6^K1425TAG^ compared to the Na_V_1.6^WT^ protein.

**Fig. 4.**
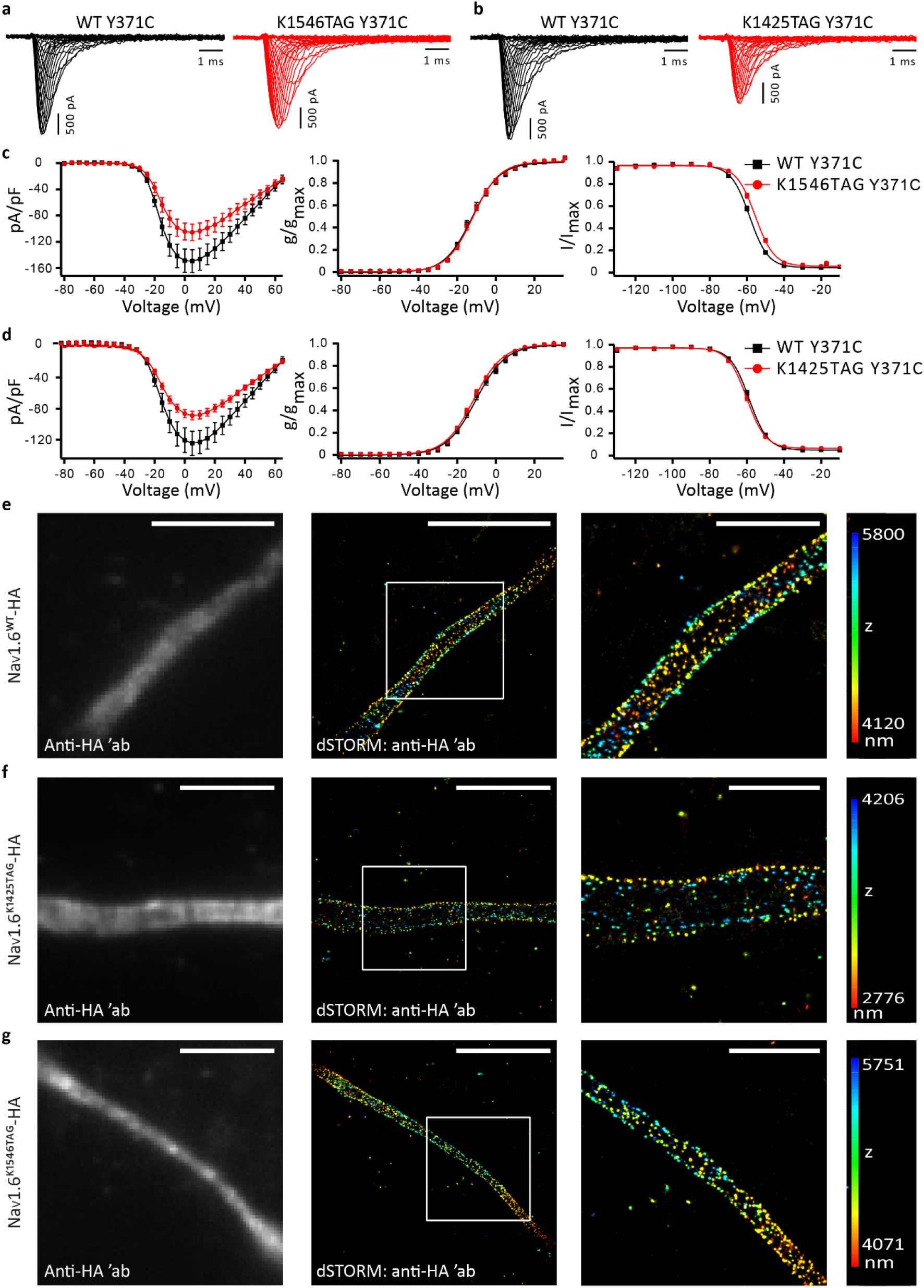
Functional characterization of clickable Na_V_1.6^TAG^-HA constructs and the effect of their expression on the AIS nanoscale organization. **a**: Representative Na^+^ current traces obtained from N1E-115-1 cells expressing either Na_V_1.6^WT, Y371C^-HA (black) or Na_V_1.6^K1546TAG, Y371C^-HA (red). **b**: Representative Na^+^ current traces obtained from N1E-115-1^β1β2^ stable cells expressing either Na_V_1.6^WT, Y371C^-P2A-eGFP (black) or Na_V_1.6^K1425TAG, Y371C^-P2A-eGFP (red). **c:** Peak Na^+^ current densities as normalized to cell capacitance, were plotted vs. voltage (left panel; number of analyzed (n) cells: n_WT_ = 20 and n_K1546TAG_ = 18), voltage-dependence of activation (middle panel; n_WT_ = 20 and n_K1546TAG_ = 18), and voltage-dependence of fast inactivation (right panel; n_WT_ = 20 and n_K1546TAG_=17) for K1546TAG (red) vs. WT comparison (black). **d**: Peak Na^+^ current densities as normalized to cell capacitance, were plotted vs. voltage (left panel; n_WT_ = 18 and n_K1425TAG_ = 20), voltage-dependence of activation (middle panel; n_WT_ = 18 and n_K1425TAG_=20), and voltage-dependence of fast inactivation (right panel; n_WT_ = 18 and n_K1425TAG_=20) for K1425TAG (red) vs. WT comparison (black). The lines represent the Boltzmann functions fit to the data points. Shown are mean ± standard errors of the mean (SEMs; **c**, **d**). Detailed statistical analyses are provided in **Supplementary Table 3**. **e**–**g**: Representative 3D direct stochastic optical (dSTORM) super-resolution images of primary rat cortical neurons at DIV 12-14 expressing NES PylRS^AF^/tRNA^Pyl^ and (**e**) Na_V_1.6^WT^-HA, (**f**) Na_V_1.6^K1425TAG^-HA, or (**g**) Na_V_1.6^K1546TAG^-HA in the presence of UAA TCO*A-Lys. Four to six days after the transfection, the neurons were click-labeled, fixed and immunostained with an anti-HA primary antibody and an Alexa Fluor Plus 647-conjugated secondary antibody. The left panels show the TIRF/HILO images of the anti-HA channel, acquired with 647 nm laser illumination prior to the 3D dSTORM imaging. The middle panels show the corresponding 3D dSTORM images, including a magnified view of the boxed region. The z positions in the 3D dSTORM images are color-coded according to the height maps shown on the right. The height maps contain minimal and maximal z position values. Scale bars: 5 μm for the TIRF/HILO and dSTORM images and 2 μm for the magnified views of the boxed regions (**e–g**).

To confirm that the nanoscale periodic organization of the Na_V_1.6 in the AIS was not affected by the overexpression of our clickable constructs or by the click labeling itself, we next performed direct stochastic optical reconstruction super-resolution microscopy (dSTORM) on the click-labeled and immunostained neurons that expressed Na_V_1.6^WT^-HA or Na_V_1.6^TAG^-HA variants. For the immunostaining, we used anti-HA primary antibody and dSTORM compatible Alexa Fluor (AF) Plus 647-conjugated secondary antibody. Immunostaining of the HA tag fused to WT or clickable Na_V_1.6 channels allowed us to directly compare them and investigate the effect of TCO*A-Lys incorporation and click labeling on the AIS structure. As there was no obvious difference in the nanoscale organization of the Na_V_1.6^WT^-HA or Na_V_1.6^TAG^-HA channels (**Fig. 4. e-g**), these experiments confirmed that the Na_V_1.6 overexpression, TCO*A-Lys incorporation, and click labeling did not affect the nanoscale periodic organization of the sodium channels in the AIS.

### Click labeling allows to study the localization of the epilepsy-causing Na_V_1.6 variants with a loss-of-function effect in living primary mouse hippocampal neurons

We next examined whether our click labeling approach in living primary neurons can be used to study the localization of two variants of the human *SCN8A* gene that cause a generalized epilepsy (T1787P and I1654N). We have recently reported that these two variants strongly reduced the Na^+^ current density in ND7/23 cells and reduced firing in mouse hippocampal neuronal cultures compared to the WT, indicating a loss-of-function (LOF) effect (Johannesen et al., 2021). However, due to the lack of suitable labeling approaches, it has remained unclear whether these variants affect the Na_V_1.6 channel function or trafficking (thereby leading to a reduced Na^+^ current density). To answer these questions, we introduced the corresponding LOF variants (T1785P and I1652N) in our clickable mouse constructs (mNa_V_1.6^TAG^). When we transfected N1E-115-1 cells with TTX-resistant versions of the clickable mNa_V_1.6^TAG, Y371C^ channels or the mNa_V_1.6^TAG, Y371C^ LOF variants, our results were comparable to those for hNa_V_1.6 (Johannesen et al., 2021), as both LOF variants showed strongly reduced Na^+^ currents compared to the clickable mNa_V_1.6^K1425TAG^ or mNa_V_1.6^K1546TAG^ controls (**Fig. 5a-b**). Hence, we verified that the effect of the hNa_V_1.6 LOF variants was the same for the mNa_V_1.6 that we had generated. These constructs were subsequently transfected into primary neurons and click-labeled to visualize the membrane population of the Na_V_1.6 channels. Since the LOF variants were previously studied in mouse hippocampal neurons (Johannesen et al., 2021), we also used this neuron type for the localization study. We observed that both LOF variants could be labeled extracellularly with click chemistry, suggesting that the channels are expressed in the AIS (**Fig. 5c** shows representative images for the K1546TAG control and LOF clickable variants). Our quantitative analysis revealed that the AIS fluorescence intensities of both LOF Na_V_1.6 variants did not significantly differ from that of the control (**Fig. 5d, Supplementary Table 4**). Therefore, our data suggest that the two variants causing reduced Na^+^ current density affect the channel conductance or opening probability rather than its trafficking to the membrane.

**Fig. 5.**
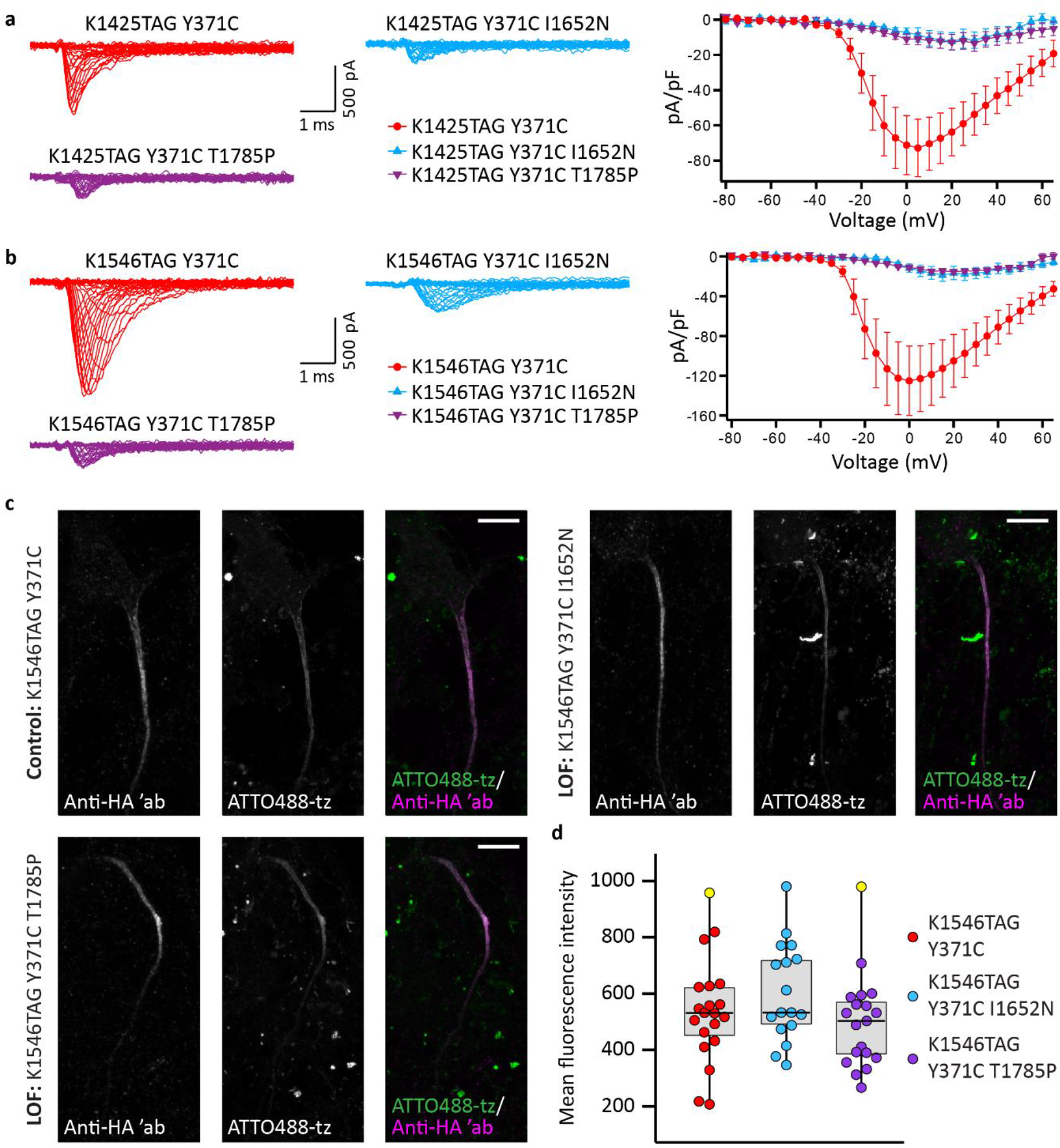
Click labeling allows localization study of the epilepsy-causing Na_V_1.6 variants with loss-of-function effect (LOF). **a**–**b**: Na^+^ current recordings of mouse Na_V_1.6 pathogenic variants combined with TAG mutations in N1E-115-1 cells. **a**: Representative Na^+^ current traces obtained from N1E-115-1 cells expressing Na_V_1.6^K1425TAG, Y371C^-HA (red), Na_V_1.6 ^K1425TAG, Y371C, I1652N^-HA (blue), or Na_V_1.6 ^K1425TAG, Y371C, T1785P^-HA (violet). Combined with the K1425TAG Y371C mutant, both the I1652N and T1785P variants significantly reduced the peak Na^+^ current density compared to the K1425TAG Y371C mutant alone (Na_V_1.6^K1425TAG, Y371C^-HA: −72.9 ± 16.4 pA/pF, number of analyzed cells (n) = 11; Na_V_1.6^K1425TAG, Y371C, I1652N^-HA: −12.3 ± 5.0 pA/pF, n = 8, p = 0.0015; Na_V_1.6^K1425TAG, Y371C, T1785P^-HA: −12.7 ± 4.4 pA/pF, n = 12, p = 0.0006; ANOVA on ranks with Dunn’s *posthoc* test). **b**: Representative Na^+^ current traces obtained from the N1E-115-1 cells expressing Na_V_1.6^K1546TAG, Y371C^-HA (red), Na_V_1.6^K1546TAG, Y371C, I1652N^-HA (blue), or Na_V_1.6^K1546TAG, Y371C, T1785P^-HA (violet). When combined with the K1546TAG Y371C mutant, both the I1652N and T1785P variants significantly reduced the peak Na^+^ current density compared to the K1546TAG Y371C mutant alone (Na_V_1.6^K1546TAG, Y371C^-HA: −125.1 ± 34.8 pA/pF, n = 13; Na_V_1.6^K1546TAG, Y371C, I1652N^-HA: −18.7 ± 6.3 pA/pF, n = 8, p = 0.0018; Na_V_1.6^K1546TAG, Y371C, T1785P^-HA: −15.4 ± 2.0 pA/pF, n = 8, p=0.0014; ANOVA on ranks with Dunn’s *posthoc* test). Shown are the mean ± standard errors of the mean (SEMs; **a**, **b**). **c**: Representative confocal images of mouse hippocampal neurons at DIV 12 expressing NES PylRS^AF^/tRNA^Pyl^, Na_V_1.6^K1546TAG, Y371C^-HA, Na_V_1.6^K1546TAG, Y371C, I1652N^-HA, or Na_V_1.6^K1546TAG, Y371C, T1785P^-HA in the presence of the TCO*A-Lys. Four days after the transfection, the neurons were click-labeled with ATTO488-tetrazine(tz), fixed, and immunostained with an anti-HA primary antibody and an Alexa Fluor 555-conjugated secondary antibody. **d**: Distribution of the mean ATTO488-tz fluorescence intensity measured in confocal images of click-labeled mouse neurons expressing the control (Na_V_1.6^K1546TAG, Y371C^-HA) or one of the LOF variants (Na_V_1.6^K1546TAG, Y371C, I1652N^-HA or Na_V_1.6^K1546TAG, Y371C, T1785P^-HA). The box plots indicate the median (the black lines inside the box), the 25^th^ and 75^th^ percentiles (the box boundaries), the single data points (the dots), and the outliers (the yellow dots). The whiskers lengths are defined by the minimum and maximum data points. The non-parametric Kruskal-Wallis test did not show any significant differences between the control and the LOFs (p = 0.209, n_K1546TAG Y371C_ = 20, n_K1546TAG Y371C I1652N_ = 17 and n_K1546TAG Y371C T1785P_ = 19). Detailed statistical analyses are provided in **Supplementary Table 4**. The Z-stack images are shown as maximum intensity projections. Scale bars: 10 μm (**c**).

### Adeno-associated virus (AAV)-based vectors for delivery of orthogonal translational machinery to primary neurons

The efficiency of transient transfection in terminally differentiated cells such as primary neurons is generally low. This is especially a problem for transfections of multiple plasmids and large genes, such as those used for click labeling of Na_V_1.6. To overcome this limitation, we developed AAV vectors as tools for delivering NESPylRS^AF^ and tRNA^Pyl^ to primary neurons. AAVs had been used in a proof-of-concept study that showed amber codon suppression of a fluorescent reporter protein in mouse neurons (Ernst et al., 2016).

To find a suitable AAV capsid for neuron transduction, we first probed the engineered variants AAV9A2 and AAV7A2. In previous work, we had generated these variants by inserting the small peptide NYSRGVD (called A2) into exposed regions on the capsids of the AAV serotypes AAV7 or AAV9, respectively, and we found that they efficiently transduced a wide variety of cell types (Borner et al., 2020). In this study, we transduced primary neurons with AAV9A2 or AAV7A2 that bore a fluorescent reporter which consisted of the nuclear localization signal (NLS)-mCherry and GFP^Y39TAG^ fusion. We observed that the neurons that had been transduced with the AAV9A2 variant started producing mCherry earlier and at slightly higher levels than the neurons that had been transduced with the AAV7A2. We then co-transduced neurons with AAV9A2 that carried the NLS-mCherry-GFP^Y39TAG^ fluorescent reporter and with AAV9A2 that carried different combinations of promoters and genes for UAA incorporation (**Supplementary Fig. 8a**). Tested promoters and genes included multiple copies of tRNA^Pyl^ and the mutant eRF1^E55D^ elongation factor (Schmied et al., 2014) which had been used to increase the efficiency of amber codon suppression, as well as conventional and minimal CMV promoters for the expression of NESPylRS^AF^ and eRF1^E55D^. These experiments revealed that a high number of neurons expressed both NLS-mCherry and GFP^Y39TAG^ in all the tested conditions (**Supplementary Fig. 8b**), which confirmed that our AAVs can be used for successful amber codon suppression of a reporter fluorescent protein. Subsequently, we tried to use these AAV vectors for click labeling of Na_V_1.6. Since the AAVs have a limited packaging capacity, we were not able to pack large *mSCN8A* genes into an AAV. Instead, for click labeling of Na_V_1.6, we combined transfection of the clickable Na_V_1.6^TAG^ plasmids and transduction with the AAV9A2 vectors that carried components for UAA incorporation. Similar to the experiments with the mCherry-GFP^Y39TAG^ reporter (**Supplementary Fig. 8**), we transduced neurons with combinations of AAV9A2 vectors that carried different orthogonal translational machinery genes. Although all the tested combinations resulted in successful click labeling (data not shown), a combination of AAVs that bore NES PylRS^AF^ and four copies of tRNA^Pyl^ showed the lowest background (**Fig. 6**). In addition, the microscopy experiments revealed that the expression level (**Fig. 6b**) and the number of neurons that expressed clickable Na_V_1.6 were higher than in the case of the transfections (50 transduced neurons vs. ^~^5-15 transfected per well of an eight-well Lab-Tek II chambered cover glass). In conclusion, we developed AAV viral vectors that enabled successful genetic code expansion and click labeling in primary neurons.

**Fig. 6.**
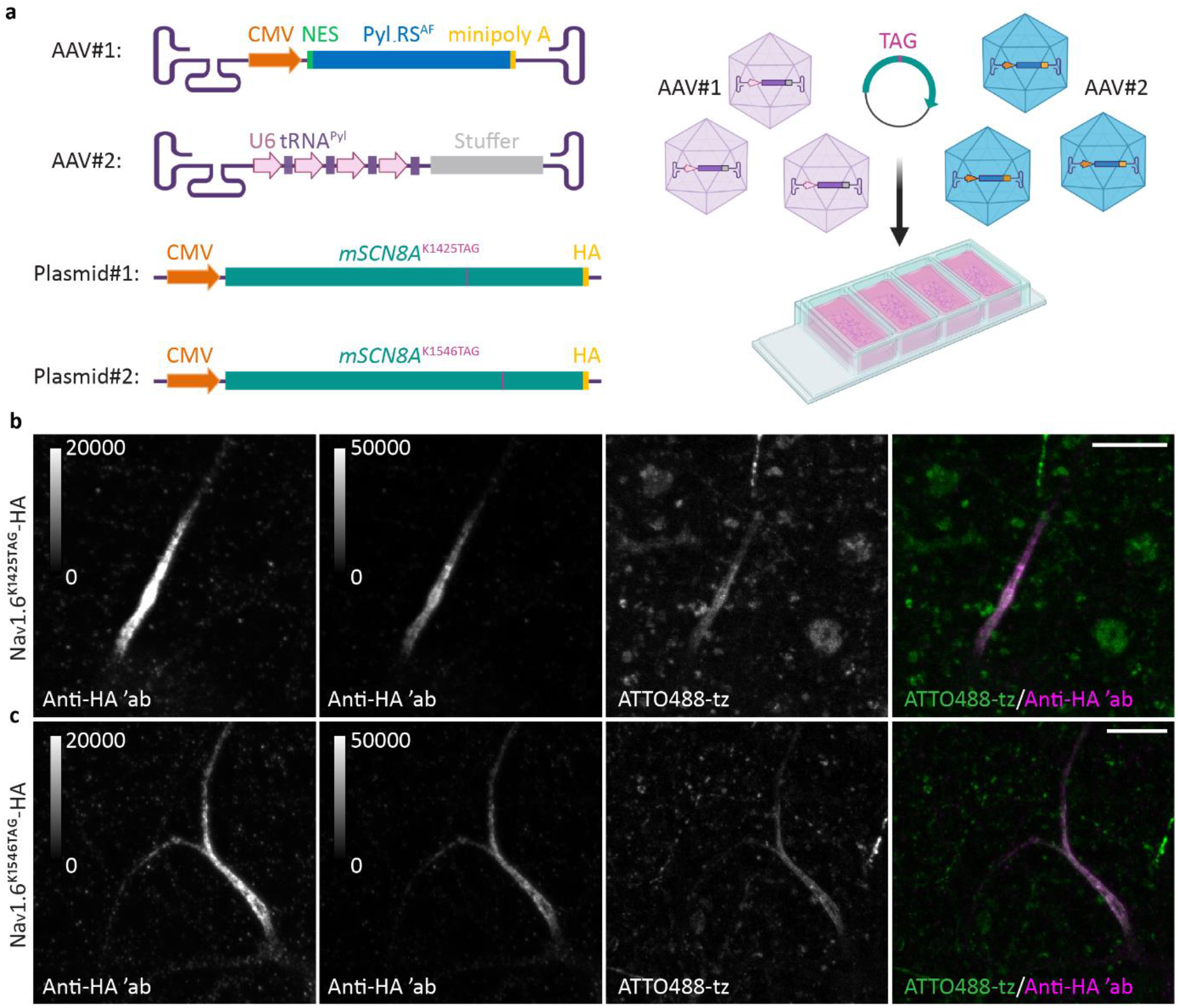
Adeno-associated virus (AAV)-based vectors are an efficient tool for delivery of orthogonal translational machinery for click labeling of Nav1.6 in primary neurons. **a:** Schematic representation of a combination of plasmid transfection and transduction with AAV9A2 viral vectors in primary rat cortical neurons. **b**–**c**: Representative confocal images of neurons at DIV 11 expressing NES PylRS^AF^/4xtRNA^Pyl^ and (**b**) Na_V_1.6^K1425TAG^-HA or (**c**) Na_V_1.6^K1546TAG^-HA in the presence of UAA TCOA*-Lys. The neurons were transfected with clickable Na_V_1.6 and transduced with orthogonal translational machinery components: NES PylRS^AF^ (AAV#1) and four copies of tRNA^Pyl^ (AAV#2) at DIV 8. Prior to imaging, living neurons were click-labeled with ATTO488-tetrazine (tz), fixed, and immunostained with an anti-HA primary antibody, followed by an Alexa Fluor 555 secondary antibody. The Z-stack images are shown as maximum intensity projections. For comparison with the transfected neurons (as shown in Fig. 3c), the brightness and contrast of the panels showing the HA channel were linearly adjusted to show the same display range (0–20,000). In addition, they were adjusted to show a broader display range (0–50,000), as indicated by the look-up-table (LUT) intensity scale bar. LUT intensity scale bars show the minimum and maximum grey values. Scale bars: 10μm (**b**–**c**).

## Discussion

Since the AIS organization is highly complex and unique, care should be taken when choosing a live-cell labeling method for microscopy studies of its components. In this study, we combined genetic code expansion and bioorthogonal click chemistry to perform direct fluorescent labeling of two AIS components: the cell adhesion molecule NF186 and the voltage-gated ion channel Na_V_1.6. The main advantage of our labeling approach is that the small fluorescent dye is directly attached to the proteins of interest in living neurons. Importantly, a clickable UAA in a complex with a dye is much smaller (^~^0.5–2 kDa) than fluorescent proteins (^~^30 kDa) and antibodies (over 150 kDa). Therefore, the protein of interest is modified in a minimally invasive way. This is of particular importance for the labeling of Na_V_, where a single mutation can severely impair the function of the channel (Johannesen et al., 2021, Meisler et al., 2021, Sole and Tamkun, 2020). Since the UAAs are introduced site-specifically into the protein of interest, positions and domains that are important for the function of the target protein can be avoided. For these reasons, click chemistry represents a powerful approach for live-cell labeling of AIS components.

Although click labeling has been used for other smaller neuronal proteins and cytoskeleton (Arsić et al., 2022, Bessa-Neto et al., 2021), NF186 and Na_V_1.6 are challenging targets due to their large size and spatially restricted expression at the AIS. For technical reasons, we first established click labeling of the smaller AIS component, NF186 (186 kDa). Although we showed that several clickable variants were suitable for fluorescent labeling, we continued using the K680TAG variant due to its bright labeling. Labeling of Na_V_1.6 was more challenging due to its complex structure and larger size (260 kDa). We unsuccessfully attempted click labeling in two host cell lines before we were able to establish it in primary neurons. For this purpose, we probed two positions in extracellular loops of Na_V_1.6: K1546 and K1425. It was reported previously that incorporation of the 17-amino-acid-long BAD domain at position K1546 minimally affected the function of Na_V_1.6 (Akin et al., 2015). Akin et al. showed that the current density measured in rat hippocampal neurons was reduced, while other biophysical parameters measured in ND7/23 cells were not affected. In this study, we observed that both clickable variants reduced the current density, although this reduction appeared statistically significant only in the case of Na_V_1.6^K1425TAG^. The reduction in the current density of clickable Na_V_1.6 is not surprising due to the fact that suppression of amber codons is not 100% efficient (Arsić et al., 2022, Bartoschek et al., 2021, Nikic-Spiegel, 2020). In addition, we observed a small shift of 2.8 mV in the inactivation curve of the clickable Na_V_1.6^K1546TAG^ variant, while the Na_V_1.6^K1425TAG^ clickable variant did not alter the biophysical properties of the channel. However, the expression and click labeling efficiency of K1425TAG were lower than those of K1546TAG. Hence, we tried to increase the expression level of K1425TAG, first by changing the transfection reagent and later by using AAV-based viral vectors to deliver orthogonal translational machinery. However, even with higher expression, click labeling of K1425TAG was weaker than click labeling of K1546TAG, most likely due to the reduced accessibility of the UAA for tetrazine-dye. That is why we continued to use K1546TAG mutant. Even though AAV vectors increased the expression of our clickable Na_V_1.6 constructs, it is important to address their limitations. Due to the AAV’s limited packaging capacity of ^~^5 kb, we used multiple AAVs to deliver different components of the orthogonal translational machinery. We are currently attempting to overcome this limitation by testing AAV variants with minimal promoters that allow packaging of all necessary components in one AAV. Furthermore, we were not able to pack the large *mSCN8A* gene into an AAV. To avoid the combination of plasmid transfection and AAV transduction, baculoviruses that have a high packaging capacity could be used instead. They were used previously for genetic code expansion of a fluorescent reporter protein in cells and *ex vivo* mouse brain slice culture (Chatterjee et al., 2013, Zheng et al., 2017).

In summary, we developed a minimally invasive approach for labeling of NF186, as well as for WT and epilepsy-causing pathogenic Na_V_1.6 channels, with small fluorescent dyes in living neurons. UAA-based minimal tags offer the opportunity to study localization and trafficking of NF186 and Na_V_1.6 in developing, mature, healthy, or injured neurons. Thanks to its compatibility with live-cell imaging, this labeling approach will provide new insights into the dynamics and plasticity of these proteins. Furthermore, the combination of two different tetrazine dyes (Arsić et al., 2022) or two different click reactions (Nikic et al., 2014) in a pulse-chase manner can be used to study different populations of NF186 or sodium channels. Moreover, in addition to conventional imaging, the small size of the labeling tag and the variety of available tetrazine dyes make click labeling particularly suitable for super-resolution imaging such as dSTORM or stimulated emission depletion (STED) microscopy (Arsić et al., 2022, Bessa-Neto et al., 2021). We also showed that our labeling approach can be used not only for different AIS components but also across different neuronal types (i.e., mouse hippocampal neurons and rat cortical neurons), thereby strengthening the potential of this method. Furthermore, click chemistry labeling can be easily adjusted for other neurofascin and voltage-gated sodium channel isoforms, including different disease-associated variants. This method can also be established for click labeling of other AIS and nodes of Ranvier proteins, including ion channels, for which antibodies, fluorescent protein fusions, and other labeling tags cannot be used. Finally, we developed AAV-based viral vectors to more efficiently deliver the components required for genetic code expansion to primary neurons. Together with the availability of natural or synthetic AAV capsid variants as well as hybrid parvoviral vectors with different cell type-specificities (Fakhiri and Grimm, 2021, Grimm and Zolotukhin, 2015), this will facilitate application of click chemistry-based protein engineering in more complex biological systems, such as organotypic slice cultures, organoids and animal models.

## Acknowledgments

We would like to thank Katja Widmaier for her excellent technical assistance and all the members of the Nikić-Spiegel group for their support, Dr. Rainer Spiegel for his advice on statistical analysis, the laboratories of Dr. Vann Bennett, Dr. Michael Davidson, and Dr. Rosalyn Adam for sharing plasmids that were obtained through Addgene, and the laboratory of Dr. Matthew Grubb and Dr. Christophe Leterrier for sharing MATLAB scripts and Fiji/ImageJ macros sets for AIS quantification. We would also like to thank Dr. Edvard Lemke for the gift of NES PylRS^AF^/tRNA^Pyl^, eRF1^E55D^, and (NLS)-mCherry-GFP^Y39TAG^ plasmids and Dr. Jason Chin for the gift of the plasmid that contains 4xtRNA^Pyl^ cassette. We are also grateful to George Philippos for his help with the mutagenesis of NF186-HA. This study was supported by the Emmy Noether Programme (project number 317530061 to I.N.-S.) of the German 1061 Research Foundation (DFG) and the Werner Reichardt Centre for Integrative Neuroscience (Ministry of Science Baden-Württemberg and former Excellence Cluster EXC307 from the DFG). This work was moreover enabled by funding from the DFG Collaborative Research Center SFB1129 (project number 240245660 to D.G.) and the Cluster of Excellence CellNetworks (EXC81 to D.G.). The electrophysiological experiments were supported by the Research Unit FOR-2715 of the DFG (grant Le1030/15-2 to H.L.).

## Author contributions

N.S. designed and performed the experiments that involved molecular biology, transfection of neuronal cells and primary neurons, click labeling, and microscopy. N.S. and I.N.-S. analyzed the click labeling and microscopy data. Y.L. and H.L. designed, performed, and analyzed the experiments that involved electrophysiological recordings of Na_V_1.6. A.A. optimized the conditions for the transfections and the amber codon suppression in the primary neurons and contributed to the data analysis. N.M. and D.G. designed and provided the AAV viral vectors. H.Lyu. and N.Z. performed the electrophysiological recordings of the loss-of-function Na_V_1.6 variants in the N1E-115-1 cells. N.S. and I.N.-S. prepared the figures (with the help of Y.L.) and wrote the manuscript with the inputs of all the authors. I.N.-S. conceived and supervised the project. All the authors reviewed and approved the final manuscript.

## Materials and Methods

### Plasmids, cloning, and mutagenesis

For click labeling of NF186, we used a plasmid that contained a rat *Nfasc* gene with a hemagglutinin (HA) tag at the C terminus. This construct was generated from a plasmid that contained a wild-type (WT) rat *Nfasc* gene expressed from the cytomegalovirus (CMV) promoter (a gift from Vann Bennett, Addgene plasmid # 31061; http://n2t.net/addgene:31061; RRID: Addgene_31061) (Zhang et al., 1998) by moving the HA tag from the N terminus to the C terminus. To delete the HA tag from the N terminus, we used the Quick Change II XL site-directed mutagenesis kit (Agilent Technologies, cat. no. 200522) following the manufacturer’s instructions. In the resulting construct, the HA tag was added to the C terminus via polymerase chain reaction (PCR)-mediated cloning using the ApaI and NotI restriction sites (resulting plasmid: CMV-NF186^WT^-HA). Clickable NF186-HA mutants (CMV-NF186^TAG^-HA) were generated by introducing amber stop (TAG) codons in the *Nfasc* gene of the original Addgene plasmid at positions K534 and K680, or by modifying CMV-NF186^WT^-HA by introducing TAG codons in the *Nfasc* gene at positions K519, K571, K604, or K809. All modifications were introduced by PCR-based site-directed mutagenesis. In the final experiments, CMV-NF186^WT/TAG^-HA was used. For click labeling of NF186 in primary neurons, the CMV promotor was excised from CMV-NF186^WT^-HA and six CMV-NF186^TAG^-HA plasmids and replaced with the human neuron-specific enolase 2 promotor (hNSE) by using the AseI and BgIII restriction sites. The hNSE promoter was amplified from the pGL3 NSE plasmid (a gift from Rosalyn Adam, Addgene plasmid # 11606; http://n2t.net/addgene:11606; RRID: Addgene_11606) (Kim et al., 2004). We used One Shot^™^ TOP10 Electrocompetent *E. coli* (Thermo Fisher Scientific, cat. no. C40452) in all the experiments that involved mutagenesis, cloning, and amplification of the NF186 plasmids, except for the experiment that involved the deletion of the HA tag, where we used the XL 10-Gold Ultracompetent Cells (Agilent, cat. no. 200315) provided with the mutagenesis kit.

Mouse 654 base pair-long *SCN1B* (*mSCN1B*, Clone ID: OMu07915D ORF clone, accession no. NM:011322.2) and mouse 558 base pair-long *SCN2B* (*mSCN2B*, clone ID: Omu42415D ORF clone, accession no. XM:006510629.3 ORF sequence) cDNAs cloned into the pcDNA3.1+/C-(k)-DYK vectors were obtained from GenScript. For the electrophysiological recordings of the Na^+^ currents, we made a multigene plasmid that contained monomeric enhanced-GFP (mEGFP) and the *mSCN1B* and *mSCN2B* genes using the MultiBacMam system^™^ kit (Geneva Biotech). We first cloned *mSCN1B* into the pACEMam2 acceptor vector (pACE Mam2 β1) and *mSCN2B* into the pMDS donor vector (pMDS β2) using the NheI and KpnI restriction sites, and then we cloned mEGFP into the pMDC donor vector (pMDC mEGFP) using the BamHI and Xbal restriction sites. The donor and acceptor vectors were components of the MultiBacMam system^™^ kit, while cDNA encoding for mEGFP was amplified from the mEGFP-N1 plasmid (a gift from Michael Davidson, Addgene plasmid #54767; http://n2t.net/addgene:54767; RRID: Addgene_54767). The multigene construct (pACEMam2_mβ1_mβ1_mEGFP) was made via CreLox recombination between donor vectors and acceptor vector, following the manufacturer’s instructions. The donor vectors were propagated in pirHC cells (Geneva Biotech), while all the other plasmids were propagated in One Shot^™^ TOP10 Electrocompetent^™^ *E. coli*. For the generation of the N1E-115^β1β2^ stable cell lines, we cloned m*SCN1B* or m*SCN2B* into the PiggyBAC-CMV-MCS-EF1-Puro cDNA/miRNA Cloning and Expression vector (BioCat, cat. no. PB510B-1-SBI) using the NheI and NotI restriction sites (PiggyBAC mβ1, PiggyBAC mβ2).

Mouse voltage-gated sodium channel 5934 base pair-long genes (*mSCN8A*^WT^, *mSCN8A*^K1425TAG^, and *mSCN8A*^K1546TAG^ were synthetized by GenScript (*mSCN8A*; transcript variant 1, NCBI Reference Sequence: NM_001077499.2) and cloned into pcDNA3.1-P2A-eGFP vectors using EcoRI and Xbal restriction sites. The final plasmids contained the *mSCN8A* gene, followed by the Xbal restriction site, self-cleaving (22 amino acids) P2A sequence, eGFP, and TAA stop codon (Na_V_1.6-P2A-eGFP). For the establishment of Na_V_1.6 click labeling, P2A-eGFP was excised and replaced with the HA tag using the Apal and XbalI restriction sites (**Supplementary Table 5**). In the final plasmids, the open-reading frame (ORF) contained the *mSCN8A* gene, followed by the HA tag and a TAA stop codon (Na_V_1.6-HA). The HA tag oligonucleotide strand was synthetized by Sigma-Aldrich as two complementary single-stranded oligonucleotides (**Supplementary Table 5**). For the electrophysiological recordings of the Na^+^ currents, the Na_V_1.6-P2A-eGFP and Na_V_1.6-HA plasmids were rendered TTX-resistant by introducing the previously described Y371C point mutation into the *mSCN8A*^WT^ and *mSCN8A*^TAG^ genes (Leffler et al., 2005, Liu et al., 2019). To study the localization of the LOF mNa_V_1.6-HA variants, we introduced I1652N or T1785P mutations (Johannesen et al., 2021) in the *mSCN8A*^WT, Y371C^, *mSCN8A*^K1425TAG, Y371C^, and *mSCN8A*^K1546TAG, Y371C^ genes. Mutagenesis of *mSCN8A* was performed using the Quick Change II XL site-directed mutagenesis kit. In the experiments that involved cloning, mutagenesis, and amplification of *mSCN8A*, we used chemically-competent XL-10 Gold Ultracompetent cells. All the steps that involved propagation of Na_V_1.6 in bacteria were performed at 27—28°C (Feldman and Lossin, 2014, O’Brien and Meisler, 2013) to avoid the introduction of additional mutations and rearrangements of *mSCN8A*.

For the incorporation of TCO*A-Lys into of NF186^TAG^-HA and Na_V_1.6^TAG^-HA, we used a recently described pcDNA3.1/Zeo(+) plasmid that contained the codon-optimized *Methanosarcina mazei-derived* Y306A/Y384F (AF) double-mutant pyrrolysyl (Pyl) tRNA synthetase fused to a nuclear export signal (NES PylRS^AF^) and its cognate amber codon suppressor tRNA^Pyl^ (Arsić et al., 2022). For electrophysiological recordings of Na^+^ currents, we transfected N1E-115-1 cells with the codon-optimized NES PylRS^AF^/tRNA^Pyl^ and WT or K1546TAG Na_V_1.6 plasmids, while we transfected the N1E-115-1^β1β2^ cells with NES PylRS^AF^/tRNA^Pyl^, which was a gift from Dr. Edward Lemke (EMBL, Heidelberg, and IMB, Mainz), and with WT or the K1425TAG Na_V_1.6 plasmids.

All the modifications introduced into the abovementioned plasmids were confirmed by sequencing. For modifications of Na_V_1.6, the entire *mSCN8A* ORFs were sequenced to confirm that there were no additional mutations or rearrangements before they were used for the transfections. The mutagenesis primers, cloning primers, and oligonucleotide sequences are provided in **Supplementary Table 5.**

### UAAs, tetrazine-dye derivatives, and antibodies

We used the trans-cyclooct-2-en-L-lysine (TCO*A-Lys; Sirius Fine Chemicals, SICHEM, cat. no. SC-8008) UAA. A 100 mM stock solution (in 0.2 M NaOH and 15% DMSO) of TCO*A-Lys was diluted 1:4 in 1 M HEPES (Thermo Fisher Scientific, cat. no. 15630056), and added to the medium at a final concentration of 250 μM. For click labeling of NF186-HA and Na_V_1.6-HA, we used ATTO488-tetrazine (ATTO488-tz; Jena Bioscience, cat. no. CLK-010-02). A 500 μM ATTO488-tz stock solution in DMSO was diluted in a warm culture medium at a final concentration of 1.5 μM, 3 μM or 5 μM.

The primary antibodies were as follows: mouse anti-HA tag (2-2.2.14) antibody (1:1000; Thermo Fisher Scientific, cat. no. 26183); rabbit anti-HA tag (C29F4) monoclonal antibody (1:1000 for Na_V_1.6 and 1:2000 for NF186; Cell Signaling, cat. no. 3724); mouse anti-ankyrin G antibody (1:50; Santa Cruz, cat. no. 12719); and mouse anti-βIII-tubulin antibody (1:1000; BioLegend, cat. no. 801202). The secondary antibodies were as follows: goat anti-rabbit AF555 (1:500; Thermo Fisher Scientific, cat. no. A21429); goat anti-rabbit AF647 Plus [1:500; AF(+)647; Thermo Fisher Scientific, cat. no. A32733], goat anti-mouse AF633 (1:500; Thermo Fisher Scientific, cat. no. A-21052); and goat anti-mouse AF(+)647 (1:500; Thermo Fisher Scientific, cat. no. A-21236).

### Cell culture

Mouse neuroblastoma x rat neuron hybrid ND7/23 cells (ECACC 92090903, Sigma Aldrich) were grown in high-glucose Dulbecco’s Modified Eagle Medium (DMEM; Thermo Fisher Scientific, cat. no. 41965062) supplemented with 10% heat-inactivated fetal bovine serum (FBS; Thermo Fisher Scientific, cat. no. 10270106), 1% penicillin-streptomycin (PS; Sigma Aldrich, cat. no. P0781), 1% sodium pyruvate (Thermo Fisher Scientific, cat. no. 11360039), and 1% L-glutamine (Thermo Fisher Scientific, cat. no. 25030024) at 37 °C and 5% CO_2_. FBS was heat-inactivated via incubation at 56 °C for 30 min. Mouse neuroblastoma N1E-115-1 cells (ECACC 08062511, Sigma Aldrich) were grown in high-glucose DMEM supplemented with 10% FBS and 1% PS at 37 °C and 5% CO2. For maintenance of the PiggyBAC N1E-115-1^β1β2^ stable cells, in addition to 10% FBS and 1% PS we supplemented the medium with 3 μg/ml puromycin (Sigma Aldrich, cat. no. P8833). For the experiments that involved transfection and electrophysiological recordings, neuronal cells were used from passage 3-15 and were passaged three times a week.

For the microscopy experiments, neuroblastoma ND7/23 or N1E-115-1 cells were seeded on four-well Lab-Tek II chambered cover glasses (German #1.5 borosilicate glass; Thermo Fisher Scientific, cat. no. 155382) at the following densities: 50,000 ND7/23 cells per well for click labeling of NF186 and 100,000 ND7/23 cells per well, or 60,000 N1E-115-1 cells per well for click labeling of Na_V_1.6. Before cell seeding, chambered cover glasses were pre-coated with 10 μg/ml Poly-D-lysine (PDL; Sigma Aldrich, cat. no. P6407) solution in double-distilled water (ddH_2_O) for at least 4 h at room temperature (RT). The chambered cover glasses were washed three times with ddH_2_O and dried completely before cell seeding. For the electrophysiological recordings, 250,000 N1E-115-1 or 160,000–200,000 PiggyBAC N1E-115-1^β1β2^ stable cells were seeded per well of a six-well plate a day before the transfection.

For the experiments that included genetic code expansion and click labeling of NF186 and Na_V_1.6, Gibco primary rat cortex neurons, from Sprague Dawley embryonic day 18 rats (Thermo Fisher Scientific, cat. no. A365512) were thawed and cultured based on the manufacturer’s recommendation. The neurons were maintained in the B-27^™^ Plus Neuronal Culture System (Thermo Fisher Scientific, cat. no. A3653401) that contained 2% B27 Plus and 1% PS. For the microscopy experiments, rat neurons were seeded in eight-well Lab-Tek II chambered cover glasses (German #1.5 borosilicate glass; Thermo Fisher Scientific, cat. no. 155409) at a density of 100,000–110,000 neurons per well. Prior to the neuron seeding, chambered cover glasses were pre-coated with 20 μg/ml of PDL solution in ddH_2_O for 2 h at RT. Afterwards, they were washed three times with ddH_2_O, dried completely, and pre-incubated with 250 μl of a warm culture medium for at least 30 min at 37 °C, 5% CO_2_. Half of the culture medium was replaced every 3–4 days.

For click labeling of the LOF Na_V_1.6 variants, we used mouse hippocampal neurons. Our animal protocols for the mouse hippocampal neuronal culture preparation were approved by the local Animal Care and Use Committee (Regierungspraesidium Tübingen, Tübingen, Germany). Neurons were isolated from embryonic day 18 C57BL/6NCrl mouse pups. The pregnant mice were cervically dislocated after asphyxiation with CO_2_, and then the embryos were taken out and decapitated immediately. The hippocampi within the whole brain stored in cold magnesium- and calcium-free Hanks’ Balanced Salt Solution (HBSS; Thermo Fisher Scientific, cat. no. 14175053) were recognized under a dissecting microscope (Olympus SZ 61, Shinjuku, Tokyo, Japan) and isolated using fine forceps. After the hippocampi were washed three times with a cold HBSS solution, the hippocampal tissue was incubated for 14 min in 2.5% trypsin (Thermo Fisher Scientific, cat. no. 15090046) at 37°C and then washed in DMEM with FBS (PAN Biotech, cat. no. 3306-P131004) to block the enzyme digestion. Dissociated neurons were obtained via gentle mechanical trituration and plated on four-well Lab-Tek II chambered cover glasses at a density of 120,000 neurons per well. The chambered glasses were pre-coated with 0.1 mg/ml of PDL solution in ddH2O prior to embryo preparation. After 6 h, during which the neurons settled on the chambered coverslips in a 5% CO_2_ humidified atmosphere at 37 °C, the culture medium was replaced by the Neurobasal culture medium (Thermo Fisher Scientific, cat. no. 21103049) supplemented with B27 (Thermo Fisher Scientific, cat. no. 17504044), L-glutamine (Thermo Fisher Scientific, cat. no. 25030024), and PS (Thermo Fisher Scientific, cat. no. 15140122). Half of the neuronal culture medium was exchanged every 3–4 days.

### Generation of stable cell lines

For the generation of N1E-115-1^β1β2^ stable cells, we used the PiggyBac Transposon system obtained from BioCat. Before we started the generation of the stable cell line, we determined that the appropriate puromycin concentration for the selection of stable clones was 3 μg/ml. N1E-115-1 cells were seeded on a six-well plate at a density of 200,000 cells per well a day before the transfection. The following day, the cells were transfected with 0.5 μg PiggyBAC mβ1 plasmid, 0.5 μg PiggyBAC mβ2 plasmid, and 0.4 μg Super PiggyBAC Transposase Expression vector (BioCat, cat. no. PB210PA-1-SBI). For the transfection, we used the JetPrime (Polyplus-transfection, cat. no. 114-15) reagent following the manufacturer’s instructions (2.8 μl of JetPrime transfection reagent per 1.4 μg of DNA). Cells were incubated with the transfection mix for 4.5 h, and afterwards, medium was replaced. The day after the transfection, the selection was initiated by adding 3 μg/ml puromycin. To ensure that all the untransfected cells were eliminated, we increased the puromycin concentration for selection from 3 μg/ml to 6 μg/ml and added it to the cells a day after the transfection, in the evening. Seventy-two hours after the transfection, the cells were transferred from six-well plates to p10 Petri dishes (Greiner Bio-one, cat. no. 664160). The cells were further propagated in p10 Petri dishes and p20 Petri dishes (Greiner Bio-one, cat. no. 639160) until single cell clusters were formed. After the cell cluster formation, we selected single clones and propagated them further in p10 dishes. Stocks of various clones were frozen for further analysis. To confirm that *mSCN1B* and *mSCN2B* were stably incorporated into the genomes of the N1E-115-1 cells, we extracted genomic(g)DNA from different clones with the PureLink gDNA kit (Thermo Fisher, cat. no. K1820-01) and PCR-amplified *mSCN1B* and *mSCN2B* genes from the gDNA. The clones with correct patterns on the gel were used in the experiments. The primers used for amplification of gDNA are given in **Supplementary Table 5.**

### Transfections

For click labeling of NF186, ND7/23 cells were transfected a day after they were seeded with the Lipofectamine 2000 transfection reagent (Thermo Fisher Scientific, cat. no. 11668027), as described in detail (Arsić et al., 2022, Arsić and Nikić-Spiegel, 2022). Briefly, the cells were seeded into four-well Lab-Tek II chambered cover glasses and transfected with a total DNA amount of 1 μg per well (0.5 μg WT/TAG plasmid and 0.5 μg NES PylRS/tRNA^Pyl^ plasmid) and a DNA/Lipofectamine 2000 reagent ratio of 1 μg/2.4 μl. After the addition of the transfection mix, TCO*A-Lys/1M HEPES was added to the medium at a final concentration of 250 μM. After 6 h, the medium was replaced, TCO*A-Lys/1M HEPES was diluted in the same way as described above and added again to the medium, and the cells were incubated overnight. The following day, the cells were click-labeled and immunostained.

For click labeling of Na_V_1.6, ND7/23 or N1E-115-1 cells were transfected a day after they were seeded with Lipofectamine 3000 transfection reagent (Thermo Fisher, cat. no. L3000015). We used a DNA/Lipofectamine 3000 reagent ratio of 1 μg/1.5 μl and a DNA/P3000 ratio of 1 μg/2 μl. For the microscopy experiments, ND7/23 or N1E-115-1 cells were seeded into four-well Lab-Tek II chambered cover glasses and transfected using a total of 1.8 μg of DNA per well (0.8 μg Na_V_1.6 WT/TAG, 0.8 μg NES PylRS/tRNA^Pyl^, 0.1 μg pACEmam2 mβ1, and 0.1 μg pMDC mβ2). For the electrophysiological recording experiments, N1E-115-1 cells were seeded into six-well plates and transfected with a total amount of 8.4 μg DNA per well (4 μg Na_V_1.6, 4 μg NES PylRS/tRNA^Pyl^, and 0.4 μg pACEMam2_mβ1_mβ2_mGFP). N1E-115-1^β1β2^ cells were seeded into six-well plates and transfected with a total amount of 5 μg of DNA per well [2.5 μg Na_V_1.6-P2A-eGFP and 2.5 μg NES PylRS/tRNA^Pyl^ (which was a gift from Dr. Edward Lemke)]. After the addition of the transfection mix, TCO*A-Lys/1 M HEPES was added to the medium at a final concentration of 250 μM. After 6 h, the medium was replaced and TCO*A-Lys was diluted in the same way as described above and then added again to the medium. After 2 days of incubation at 37°C and 5% CO_2_, the ND7/23 or N1E-115-1 cells were click-labeled for the microscopy experiments. For the whole-cell patch clamp recordings, the cells were assessed for a fluorescent signal, counted, and reseeded in 35 mm Petri dishes (Greiner Bio-one, cat. no. 627160) at a density of around 180,000 cells per dish without further addition of TCO*A-Lys/1 M HEPES. The cells were left at 37°C and 5% CO_2_ for 2–4 h until the electrophysiological recordings were performed. The cells were always recorded on the same day.

Primary Sprague Dawley rat cortical neurons were transfected with Lipofectamine 2000 as described previously (Arsić et al., 2022, Arsić and Nikić-Spiegel, 2022) or with Lipofectamine 3000 following the manufacturer’s instructions. For click labeling of NF186, neurons were transfected with Lipofectamine 2000 on DIV 7; and for click labeling of Na_V_1.6, neurons were transfected with Lipofectamine 2000 or 3000 on DIV 8. Neurons were seeded into eight-well Lab-Tek II chambered cover glasses, and a total amount of 1 μg per well was used for the transfection with Lipofectamine 2000 (0.5 μg Na_V_1.6 WT/TAG or NF186 WT/TAG and 0.5 μg NES PylRS/tRNA^Pyl^). For the transfection with Lipofectamine 3000, we used a total amount of 0.5 μg per well (at the DNA/Lipofectamine 3000 ratio of 1 μg/1.5 μl and the DNA/P3000 ratio of 1 μg/2 μl). The transfection mix was prepared in 25 μl Opti-MEM Reduced serum media (Thermo Fisher, cat. no. 31985062) without the addition of antibiotics. Then 250 μl of the medium was removed and saved for later at 37°C and 5% CO_2_ (conditioned medium). The entire transfection mix was added to the well that contained the remaining 250 μl of the medium. The neurons that were transfected with Lipofectamine 2000/3000 were incubated with the transfection mix for at least 6 h. Afterwards, the transfection mix was removed, 250 μl of the previously collected conditioned medium was added, and 250 μl of the fresh culture medium was added to each well. TCO*A-Lys/1 M HEPES was added to a final concentration of 250 μM, and the cells were incubated for 3 days at 37°C and 5% CO_2_. After 3 days of incubation, the medium that contained TCO*A-Lys was completely removed and replaced with one-half of the conditioned medium and one-half of the fresh culturing medium. In the controls that did not contain TCO*A-Lys/1 M HEPES, one-half of the medium was replaced with the fresh culture medium. Click labeling was performed on the following day. For localization of the LOF Na_V_1.6 variants, mouse hippocampal neurons were transfected with Lipofectamine 2000 as described previously on DIV 7–8. The only difference was that the neurons were seeded in four-well Lab-Tek II chambered cover glasses. Therefore, the amounts of the DNA and the Lipofectamine 2000 transfection reagent were scaled up to correspond to the size of the four-well Lab-Tek II chambered cover glasses (two times more DNA and Lipofectamine 2000 were used).

### Click labeling of NF186 and Na_V_1.6

Click labeling of ND7/23 cells expressing NF186-HA was performed a day after the transfection. Until the labeling, the cells were incubated with TCO*A-Lys. The cells were labeled with 1.5 μM ATTO488-tz diluted in a warm culture medium at 37°C and 5% CO_2_. After 10 min of incubation, the dye solution was removed, the cells were washed 1–2 times with 0.01 M phosphate-buffered saline (PBS; 137 mM NaCl, 10 mM Na_2_HPO_4_, 1.8 mM KH_2_PO_4_, 2.7 mM KCl, pH 7.4), fixed with 4% paraformaldehyde (PFA; Sigma–Aldrich, cat. no. 158127) in 0.1 M phosphate buffer (PB) for 15 min at RT and washed 3 times with PBS. Click labeling of ND7/23 or N1E-115-1 cells expressing Na_V_1.6^WT^-HA or Na_V_1.6^TAG^-HA was performed using the same procedure described previously except that click labeling was performed 40–46 h after the transfection with 3 μM ATTO488-tz.

Click labeling of living rat cortical neurons or mouse hippocampal neurons was performed 4 days after the transfection except for the experiments that involved dSTORM, where rat neurons were click-labeled 4-6 days after the transfection. The medium that contained TCO*A-Lys was removed a day before the click labeling. For the click labeling of NF186-HA, rat neurons were labeled with 5 μM ATTO488-tz diluted in a warm culture medium for 10 min at 37°C and 5% CO_2_. After 10 min of incubation with the dye, the medium was removed and the neurons were fixed with 4% electron-microscopy-grade PFA (Electron Microscopy Sciences, cat. no. 15710) diluted in a cytoskeleton-protective buffer (PEM; 80 mM PIPES, 5 mM EGTA, 2 mM MgCl_2_; pH 6.8) for 15 min at RT and washed 3 times in PBS (5 min per wash). For the live-cell imaging of NF186, after the click labeling, the cells were washed 2–3 times with the culture medium, placed in a Hibernate E medium that contained 2% B27 Plus and 1% PS, and kept in the incubator until the imaging (37°C, 5% CO_2_). Click labeling of the rat cortical neurons or mouse hippocampal neurons expressing Na_V_1.6-HA was performed with the same procedure, except that before the click labeling, the neurons were washed 3 times with Tyrode’s solution (100 mM NaCl, 5 mM KCl, 5 mM MgCl_2_, 2 mM CaCl_2_, 15 mM D-glucose, 10 mM HEPES; pH 7.4, osmolarity 243–247 mOsm) and incubated for 3 min in 1% BSA/Tyrode’s solution (Bessa-Neto et al., 2021). After 10 min of incubation with the ATTO488-tz dye, the neurons were washed 4 times with Tyrode’s solution, fixed immediately, and washed 3 times in PBS (5 min per wash). For the live-cell imaging of Na_V_1.6, upon the click labeling, the rat neurons were placed in the Hibernate E medium that contained 2% B27 Plus and 1% PS and kept in the incubator until the imaging (37°C, 5% CO_2_).

### Immunostaining

For immunostaining of neuroblastoma cells and primary neurons, all the blocking steps were performed at RT for 1 h. Incubation with primary antibodies was performed ON at 4°C while incubation with secondary antibodies was performed at RT for 1 h. After the incubation with the primary and secondary antibodies, the cells were washed 3 times with PBS (5 min per wash). Then they were kept in PBS at 4°C until the imaging.

For immunostaining of ND7/23 cells expressing NF186-HA, the cells were permeabilized with 0.2% Tween 20/PBS for 30 min. Afterwards, the cells were blocked with 5% FBS. Mouse anti-HA primary and goat anti-mouse AF(+)647 secondary antibodies were diluted in a blocking buffer. For immunostaining of rat neurons expressing NF186-HA, the neurons were permeabilized and blocked in a buffer that contained 0.2% Triton X-100 (Sigma-Aldrich, cat. no. X100), 10% goat serum (GS; Thermo Fisher, cat. no. 16210072), and 3% BSA in PBS. Rabbit anti-HA and mouse-anti ankG primary antibodies were diluted in the blocking buffer. Goat anti-rabbit AF555 and goat-anti mouse AF633 secondary antibodies were diluted in 3% BSA/PBS.

For immunostaining of Na_V_1.6-HA, neuroblastoma cells or primary neurons were permeabilized with 0.1% Triton X-100/PBS for 10 min and blocked in a buffer that contained 10% GS and 3% BSA. Rabbit anti-HA primary antibody was diluted in the blocking buffer. Goat anti-rabbit AF555 secondary antibody was diluted in 3% BSA/PBS. For the AIS quantification of the rat cortical neurons expressing Na_V_1.6-HA, in addition to the rabbit anti-HA primary antibody and the goat-anti rabbit AF555 secondary antibody, neurons were immunostained with mouse anti-ankG primary antibody and goat anti-mouse AF633 secondary antibody.

For dSTORM imaging of Na_V_1.6, rat cortical neurons were immunostained with rabbit anti-HA antibody as described above, except that instead of goat-anti rabbit AF555, goat-anti rabbit AF(+)647 secondary antibody was used.

### Adeno-associated viruses (AAVs) production, purification and titration

A fluorescent reporter and genetic code expansion components—nuclear localization sequence (NLS)-mCherry-GFP^Y39TAG^, NES PylRS^AF^, one copy tRNA^Pyl^, and eRF1^E55D^ [gifts from Dr. Edward Lemke (EMBL, Heidelberg, and IMB, Mainz)], and a cassette containing four copies of tRNA^Pyl^ [a gift from Jason Chin (MRC Laboratory of Molecular Biology, Cambridge, UK)]—were separately cloned into a self-complementary AAV vector plasmid (encoding transgenes flanked by inverted terminal repeats) backbone (**Supplementary Fig. 8a**). AAV vectors were produced via triple-transfection of the HEK293T cells with the AAV vector plasmid, the AAV helper plasmid (encoding the AAV rep and cap genes; AAV7A2 and AAV9A2 caps were used in this study) and adenoviral helper plasmid at a 1:1:1 molar ratio using a polyethylenimine (PEI MAX; Polysciences, Warrington, USA, cat.no. 24765-1) as the transfection reagent. Ten 15 cm dishes were used to produce each AAV vector. In brief, 4 million HEK293T cells per dish were seeded 2 days before the transfection. A DNA mix (44 μg plasmids in 790 μl H_2_O and 790 μl of 300 mM NaCl per dish) and a PEI mix (352 μl PEI, 438 μl H_2_O, and 790 μl of 300 mM NaCl per dish) were mixed together, vortexed thoroughly, incubated at RT for 10 min, and added to the cells dropwise. Three days after the transfection, the cells were harvested with a cell scraper and centrifuged at 800 g for 15 min. The cell pellets were resuspended in a 5 ml Benzonase buffer (150 mM NaCl, 50 mM Tris-HCl, 2 mM MgCl2; pH 8.5) and lysed via 5 freeze-thaw cycles. Lysates were sonicated for 1 min and 20 sec and incubated with 75 U/mL Benzonase (Merck, Darmstadt, Germany, cat. no. 1.01695.0001) at 37°C for 1 h. Then the lysates were centrifuged at 4000 g for 15 min 2 times to get rid of the cell debris.

The AAV vectors in the abovementioned supernatant were purified with an iodixanol (OptiPrep^™^ (Iodixanol); Progen, Heidelberg, Germany, cat. no. 1114542) gradient. Each vector was loaded into ultracentrifugation tubes (Seton Scientific, Petulama, CA, USA) through a Pasteur pipette, followed by 2 ml of 15%, 25%, 40%, and 60% iodixanol solution. The tubes were sealed and centrifuged at 50,000 rpm at 4°C for 2 h in an OptimaTM L-90K ultracentrifuge using the 70.1Ti rotor (Beckman Coulter, Brea, CA, USA). After the centrifugation, the 1 ml solution at the interface of 40% and 60% phases was collected and stored at −80°C.

The AAV titers were measured with droplet digital PCR (ddPCR). The AAV samples were diluted at 1: 10^6^. Reaction solutions (20 μl) that contained 900 nM of each primer, 25 nM of the probe, 10 μl of the 2x ddPCR Supermix for probes (no dUTP; Bio-Rad, Hercules, USA, cat. no. 1863024), and 5 μl of the diluted AAV sample were prepared for the droplet generation. The droplets were generated with a QX200 Droplet Generator (Bio-Rad, Hercules, USA), and then transferred to a 96-well PCR plate. PCR was run in a C1000 Touch Thermal Cycler (Bio-Rad, Hercules, USA), and the results were read by QX200 Droplet Reader (Bio-Rad, Hercules, USA). The primers used for ddPCR are given in **Supplementary Table 5**.

### Transduction with AAVs and transfections of primary neurons

For the transduction of the primary neurons with AAVs, we adjusted the desired the multiplicity of infection (MOI) by diluting AAV stocks in a warm fresh culture medium supplemented with 2% B27 Plus and 1% PS. Before the transduction, an equal amount of conditioned medium was added to the prewarmed fresh medium that contained the AAV.

Primary neurons were transduced at DIV 5 with AAV9A2 or AAV7A2 encoding the NLS-mCherry-GFP^Y39TAG^ expressed from the CMV promoter without the addition of TCO*A-Lys. We used an MOI of 50,000. For the genetic code expansion of GFP^Y39TAG^, we co-transduced rat primary cortex neurons at DIV 8 with AAV9A2 vectors encoding the CMV-NLS-mCherry-GFP^Y39TAG^ (AAV#6) and the following genetic code expansion components: (a) CMV-NES PylRS^AF^ (AAV#1) and four copies of tRNA^Pyl^ (AAV#2); (b) CMV-NES PylRS^AF^ (AAV#1), 4xtRNA^Pyl^ (AAV#2), and mutant eukaryotic release factor eRF1^E55D^ expressed from the CMV promoter (AAV#3); (c) CMV-NES PylRS^AF^ (AAV#1) and AAV9A2 carrying minimal (min)CMV-eRF1^E55D^ and one copy of tRNA^Pyl^ (AAV#5); (d) AAV9A2 carrying minCMV-NES PylRS^AF^ (AAV#4) and one copy of tRNA^Pyl^ and AAV9A2 carrying mineRF1^E55D^ and one copy of tRNA^Pyl^ (AAV#5; **Supplementary Fig. 8a**). An MOI of 5,000 was used for each AAV. TCO*A-Lys/1 M HEPES was added immediately upon transduction at a final concentration of 250 μM. The neurons were assessed daily for their fluorescent signal and viability. Three days after the transductions, the neurons were fixed with the 4% PFA/PEM buffer for 15 min at RT, washed 3 times in PBS (5 min per wash), and left at 4°C until the imaging.

For click labeling of Na_V_1.6, we transfected primary rat cortical neurons at DIV 8 with 0.5 μg Na_V_1.6^TAG^-HA using Lipofectamine 2000, as described in the transfection section. After 6 h, we removed the transfection mix and added 250 μl of the condition medium that was saved previously and 250 μl of the fresh medium that contained the same combinations of AAV92A used for the NLS-mCherry-GFP^Y39TAG^. For each AAV, we used an MOI of 15,000. TCO*A-Lys/1 M HEPES was added immediately upon the transduction at a final concentration of 250 μM. The neurons were assessed daily for viability. Three days after the transfection and transduction, we click-labeled the neurons with 5 μM ATTO488-tz, as described in the Click labeling section, and fixed neurons immediately with the 4% PFA/PEM buffer for 15 min at RT. The neurons were washed 3 times in PBS (5 min per wash) and immunostained afterwards, as described in the Immunostaining section.

### Fixed- and live-cell imaging

Confocal imaging was performed on an LSM 710 confocal scanning microscope (Carl Zeiss, Oberkochen, Germany) equipped with the following laser lines (nm): 405, 440, 458, 488, 514, 561, and 633. Images were acquired with an oil Plan-Apochromat 63× objective (NA 1.4) and with the following settings: a 1024 x 1024 pixel frame size, a 16-bit image depth, 2× line averaging, a 6.30 μs pixel dwell time, and 0.132 μm pixel size. We used a 488 nm laser line to excite ATTO488-tz, a 561 nm laser line to excite AF555, and a 633 nm laser line to excite AF(+)647 or AF633. The pinhole was set at 1 Airy Unit in all the channels, and the emission light was collected sequentially. Images were acquired in two channels (488 nm and 561 nm, or 488 nm, and 633 nm) or three channels (488 nm, 561 nm and 633 nm) channels, as single planes or as Z-stacks with a 0.42 μm step size.

For live-cell confocal imaging, we used a temperature module and a heating insert (PeCon, Erbach Germany) that was warmed up to 37°C. Live rat cortical neurons were imaged in the Hibernate E medium supplemented with 2% B27 Plus and 1% PS.

Widefield epifluorescence and 3D dSTORM imaging were performed on an Inverted Nikon Eclipse Ti2-E (Nikon Instruments) equipped with an XY motorized stage; a Perfect Focus System; oil-immersion objectives (Apo 60x, NA 1.4, oil and HP Apo TIRF 100×H, NA 1.49, and Oil); an N-STORM module; the following filter cubes: 488 (AHF; EX 482/18; DM R488; BA 525/45), 561 (AHF; EX 561/14; DM R561; BA 609/54), and 647 (AHF; EX 628/40; DM660; BA 692/40); Nikon Normal STORM (T660lpxr, ET705/72 m); the following laser lines (nm): 405, 488, 561 and 647; and ORCA-Flash 4.0 sCMOS camera (Hamamatsu Photonics). The setup was controlled by NIS-Elements AR software (Nikon Instruments).

Images of rat cortical neurons expressing NF186^WT^-HA were acquired in the widefield mode with an Apo 60× oil objective, 30 ms exposure time, 1024×1024 pixel frame size, 16-bit image depth, and pixel size 0.27 μm. For the widefield imaging, a fluorescent lamp (Lumencor Sola SE II) was used as a light source, with the excitation light intensity set at 20%.

### Image processing

All the images shown in the main and supplementary figures, except for the dSTORM images, were processed in ImageJ/Fiji software (Schindelin et al., 2012). Raw confocal single planes or Z-stacks and widefield single planes images were imported in ImageJ/Fiji. The brightness and contrast of the 16-bit images were linearly adjusted. The Z-stacks were converted into maximum intensity projections prior to the linear adjustment of the brightness and contrast. For presentation purposes, the all images were converted to an 8-bit depth, exported as .tiff files, and arranged into figures using Adobe Illustrator.

The schemes presented in the manuscript were made in BioRender.com.

### 3D dSTORM imaging and image processing

3D dSTORM imaging was performed on the N-STORM module of the Inverted Nikon Eclipse Ti2-E microscope. 3D dSTORM images of the rat cortical neurons expressing Na_V_1.6 were acquired with an HP Apo TIRF 100×H objective, a 647 nm laser line (LU-NV Series Laser Unit), and a Nikon Normal STORM filter cube. The emitted light was imaged with an ORCA-Flash 4.0 sCMOS camera that contained a cylindrical lens introduced in the light path (Huang et al., 2008). Prior to the 3D dSTORM imaging, the AIS of interest was assessed for click labeling. Afterwards, an image of the neuron expressing Na_V_1.6-HA was captured with a 647 nm laser (HA channel) set at 1% of the power. Afterwards, the same neuron was imaged with 3D dSTORM. The 3D dSTORM imaging was performed in the total internal reflection fluorescence (TIRF) or highly inclined and laminated sheet microscopy (HILO) mode with continuous 647 nm laser illumination set at full power (100%). The frame size was 128 × 128 pixels, and the image depth, 16-bit. For each 3D dSTORM image, 20,000–30,000 frames were acquired at 33,3 Hz. During the acquisition, 405 nm laser illumination set at 5% was used. The 3D dSTORM was calibrated previously using TetraSpeck Microspheres (Thermo Fisher, cat. no. T7279) following NIS-Elements’ instructions. The 3D dSTORM imaging was performed in the GLOX BME buffer. GLOX BME buffer, which was prepared fresh prior to use by mixing 7 μl BME, 7 μl GLOX solution [14 mg glucose oxidase (Sigma Aldrich, cat. no. G2133)], 50 μl catalase (17 mg/ml; Sigma Aldrich, cat. no. C3155), and 200 μl Buffer A (10 mM Tris; pH 8, 50 mM NaCl) with 690 μl Buffer B (50 mM Tris,pH 8, 10 mM NaCl) that contained 20% w/v glucose (Sigma Aldrich, cat. no. D9559). The buffer was prepared on ice.

The 3D dSTORM images were processed in NIS-Elements AR software. The molecule identification settings were set at the defaults for the 3D dSTORM analysis: a minimum width of 200 nm, a maximum width of 700 nm, an initial fit width of 300 nm, a maximum axial ratio of 2.5, and a maximum displacement of 1. The minimum height for the peak detection was set at 100, and the localization analysis was performed with overlapping peak algorithms and drift correction. The 3D dSTORM images were reconstructed with a Gaussian rendering size of 10 nm in NIS Elements AR, and the final 3D dSTORM images (with z position height maps) were exported as .tiff files. The Z rejected molecules were excluded from the final images. The .tiff images were imported and arranged into figures using Adobe Illustrator. Scale bars for dSTORM images and corresponding TIRF images were added in NIS-Elements AR software. Scale bars for all the other (widefield or confocal images) were added in ImageJ/Fiji.

### AIS quantification

For the quantitative measurements of the AIS length of the rat primary neurons expressing NF186 or Na_V_1.6, we used the previously described MATLAB ais.m script (Grubb, 2021, Grubb and Burrone, 2010). Images were acquired on an LSM 710 confocal scanning microscope, as described in the fixed- and live-cell imaging section. We processed the images as described in the instructions of the ais.m script. Briefly, the confocal images used for the quantitative measurements (shown in Fig. 2d and Fig. 3f) of the AIS length in the neurons expressing NF186 and Na_V_1.6 immunostained with anti ankG were processed in ImageJ/Fiji software. Raw confocal single planes or Z-stacks were imported in ImageJ/Fiji, color channels were split, the brightness and contrast of the 16-bit images were linearly adjusted, and different color channels were exported as separate 16-bit .tiff files. The Z-stacks were converted into maximum intensity projections prior to the linear adjustment of the brightness and contrast. The processed images were imported in MATLAB and analyzed with ais.m script (Grubb, 2021, Grubb and Burrone, 2010). The AIS lengths of the neurons expressing recombinant NF186 or Na_V_1.6 (measured in the ankG channel) were calculated automatically in MATLAB using the ais.m script.

The AIS fluorescence intensity of the click-labeled mouse primary hippocampal neurons expressing WT or LOF Na_V_1.6 variants (shown in Fig. 5d) were measured in ImageJ/Fiji with a previously described macros set (https://github.com/cleterrier/Measure_ROIs). Images were acquired on an LSM 710 confocal scanning microscope, as described in the fixed- and live-cell imaging section. We processed images in ImageJ/Fiji using a previously described macros set (https://github.com/cleterrier/Process_Images) and measured the fluorescence intensity along the AIS in the click channel. Briefly, the processed images were imported in ImageJ/Fiji, and the line tracings along the AIS were generated semi-automatically using the NeuronJ plugin (Meijering et al., 2004), converted into ImageJ/Fiji regions of interest (ROIs), and saved as a .zip files. Afterwards, raw 16-bit images were imported in ImageJ/Fiji together with their corresponding previously generated ROIs, the fluorescence intensity was automatically measured in the click channel, and the results were exported as .csv files. For the statistical analysis, we used a mean intensity measured along the ROI (along the AIS line tracing) corrected by subtracting the background (the “Corr Mean”). The line width of the ROI for each AIS was individually adjusted according to the width of the AIS.

All the image analyses were performed blind per experimental group. We did not image/analyze neurons overexpressing NF186 that had a clearly mislocalized HA signal present in all the processes. Due to the lower expression level of the recombinant Na_V_1.6, we did not observe any neurons overexpressing Na_V_1.6. Therefore, we imaged all the transfected cells showing click labeling.

### Electrophysiological recordings

Standard whole-cell voltage clamp recordings were performed in ND7/23, N1E-115-1, or N1E-115-1^β1β2^ cells using an Axopatch 200B amplifier, a Digitata 1440A digitizer, and Clampex 10.2 data acquisition software (Axon Instruments, Union City, CA, USA) in the presence of 500 nM tetrodotoxin (TTX). The cells were held at −100 mV. Currents were filtered at 5 kHz and digitized at 20 kHz. Borosilicate glass pipettes had a final tip resistance of 1.8–2.5 MΩ when filled with the internal recording solution. The pipette solution contained (in mM) 10 NaCl, 1 EGTA, 10 HEPES, and 140 CsF. The pH was adjusted to 7.3 with CsOH, and the osmolarity was adjusted to 290 mOsm/kg with mannitol. The bath solution contained (in mM) 140 NaCl, 3 KCl, 1 MgCl_2_, 1 CaCl_2_, 10 HEPES, 20 TEACl (tetraethylammonium chloride), 5 CsCl, and 0.1 CdCl2. The pH was adjusted to 7.3 with CsOH, and the osmolarity, to 320 mOsm/kg with mannitol.

The electrophysiology data were analyzed as previously described (Liu et al., 2019). Briefly, the activation curve (conductance–voltage relationship) was derived from the current–voltage relationship. The latter was obtained by measuring the peak current at various step depolarizations from the holding potential of −100 mV according to the following formula:

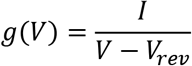

Note that g represents the conductance, I represents the recorded peak current at the test potential (V) while V_rev_ represents the apparent observed Na^+^ reversal potential.

A standard Boltzmann function was fit to the activation curves according to the following formula:

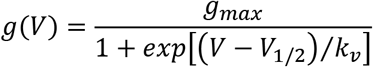

Note that g represents the conductance, I represents the recorded current amplitude at the test potential (V), V_rev_ represents the Na^+^ reversal potential, g_max_ represents the maximal conductance, V_1/2_ represents the voltage of a half-maximal activation while kV represents a slope factor.

Steady-state inactivation was determined using 100 ms conditioning pulses to various potentials followed by the test pulse to −10 mV at which the peak current reflected the percentage of non-inactivated channels. A standard Boltzmann function was fit to the inactivation curves according to the following formula:

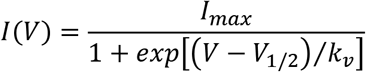

Note that I represents the recorded current amplitude at the conditioning potential (V), I_max_ represents the maximal current amplitude, V_1/2_ represents the voltage of half-maximal inactivation while k_V_ represents a slope factor.

All the data were analyzed using the Clampfit software of pClamp 10.6 (Axon Instruments), Microsoft Excel (Microsoft Corporation, Redmond, WA, USA), or Igor Pro (Wavemetrics, Portland, OR, USA).

### Statistical analyses

Statistical analyses (Shapiro-Wilk normality test, Levene’s test for equality of variances, one-way ANOVA, and Kruskal-Wallis non-parametric Hypothesis test) for the quantification of the AIS length and the quantification of the AIS fluorescence intensity were performed with IBM SPSS Statistics 28.01.0 (Armonk, New York, USA). A Shapiro-Wilk normality test indicated whether the data followed a normal distribution. The data that followed normal distribution were further tested for the homogeneity of variances. The data that met both assumptions (normality and homogeneity of variances), and did not have any significant outliers were further analyzed with one-way ANOVA with a Tukey *posthoc*. The data that followed normal distribution, but did not meet the assumption of the homogeneity of variance and/or had significant outliers were analyzed with non-parametric Kruskal-Wallis test with Dunn-Bonferroni *posthoc* (if required). The details are provided in **Supplementary Tables 1**, **2** and **4.**

The statistical analyses of the electrophysiological recordings were performed using GraphPad Prism (GraphPad Software, Inc., San Diego, CA, USA). D’Agostino & Pearson normality tests indicated whether the data followed a normal distribution. To compare the groups, the unpaired T-test was used for the data that followed a normal distribution while the non-parametric Mann-Whitney U test was used for the data that did not follow normal distribution. To compare three groups of data, ANOVA on ranks with Dunnett’s *posthoc* test was used for data that did not follow normal distribution. For all the statistical tests, the significance levels compared to the controls are indicated in the **Supplementary Table 3** using the following symbols: * p < 0.05, ** p < 0.01, and *** p < 0.001. The corresponding electrophysiology graphs were created in IgorPro (Wavemetrics, Portland, OR, USA).

