## Supplemental Data 1 for "Direct fluorescent labeling of NF186 and Na_V_1.6 in living primary neurons using bioorthogonal click chemistry"

Supplementary figures 1–8

Supplementary tables 1–5

### Supplementary figures

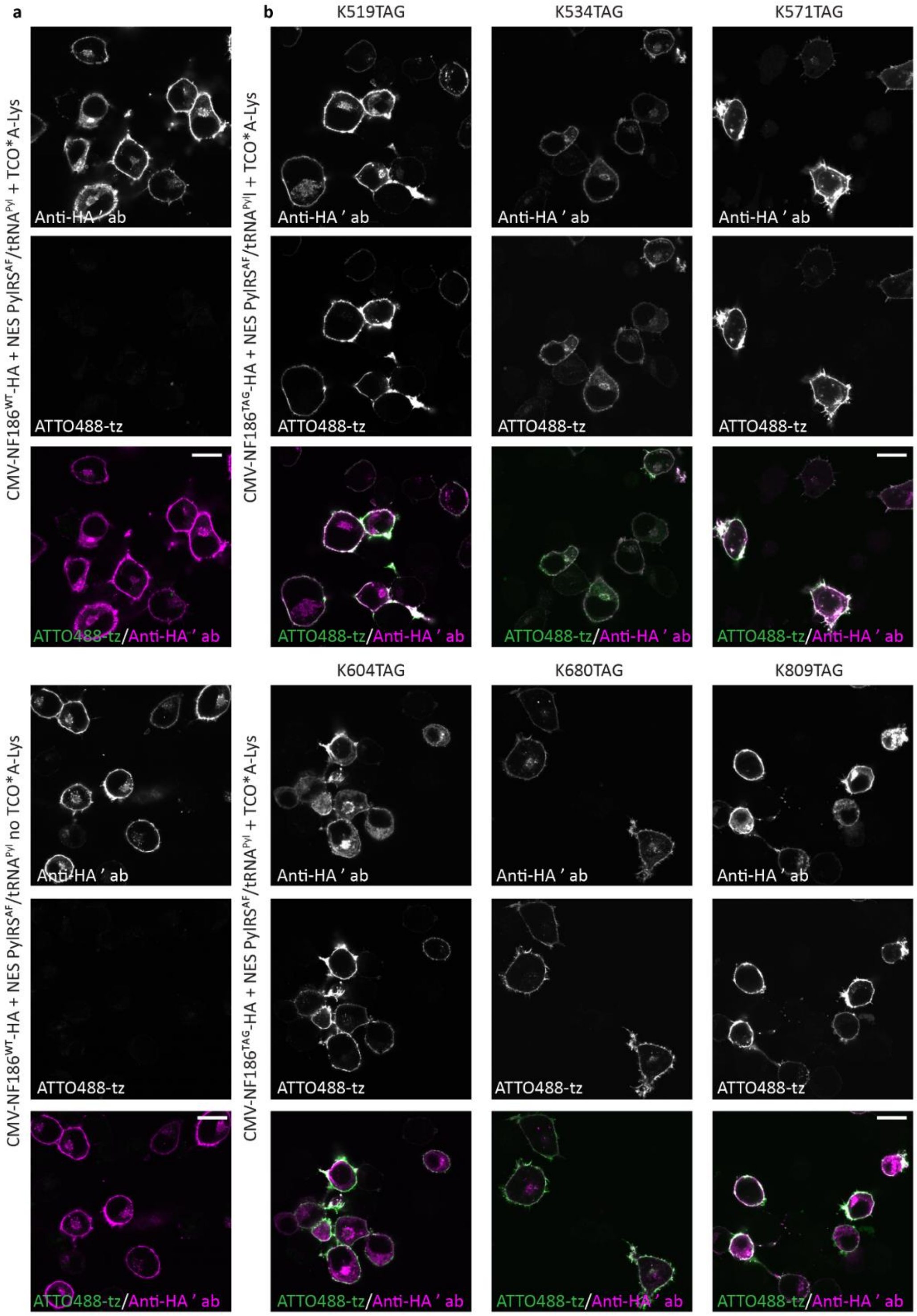


**Supplementary Fig. 1. Genetic code expansion and click labeling of NF186-HA in living ND7/23 cells.** Representative images of ND7/23 cells co-expressing NES PylRS$^{AF}$/tRNA$^{Pyl}$ and (**a**) CMV-NF186$^{WT\text{-}}$HA in presence or in the absence of the unnatural amino acid TCO*A-Lys or (**b**) one of the CMV-NF186$^{TAG}$-HA amber mutants (K519TAG, K534TAG, K571TAG, K604TAG, K680TAG, and K809TAG) in the presence of the TCO*A-Lys. A day after the transfection, the cells were click-labeled with ATTO488-tetrazine (tz), fixed, and immunostained with anti-HA primary and Alexa Fluor Plus 647-conjugated secondary antibodies. Single-plane images were acquired with a confocal scanning microscope. Scale bars: 20 µm (**a**–**b**).

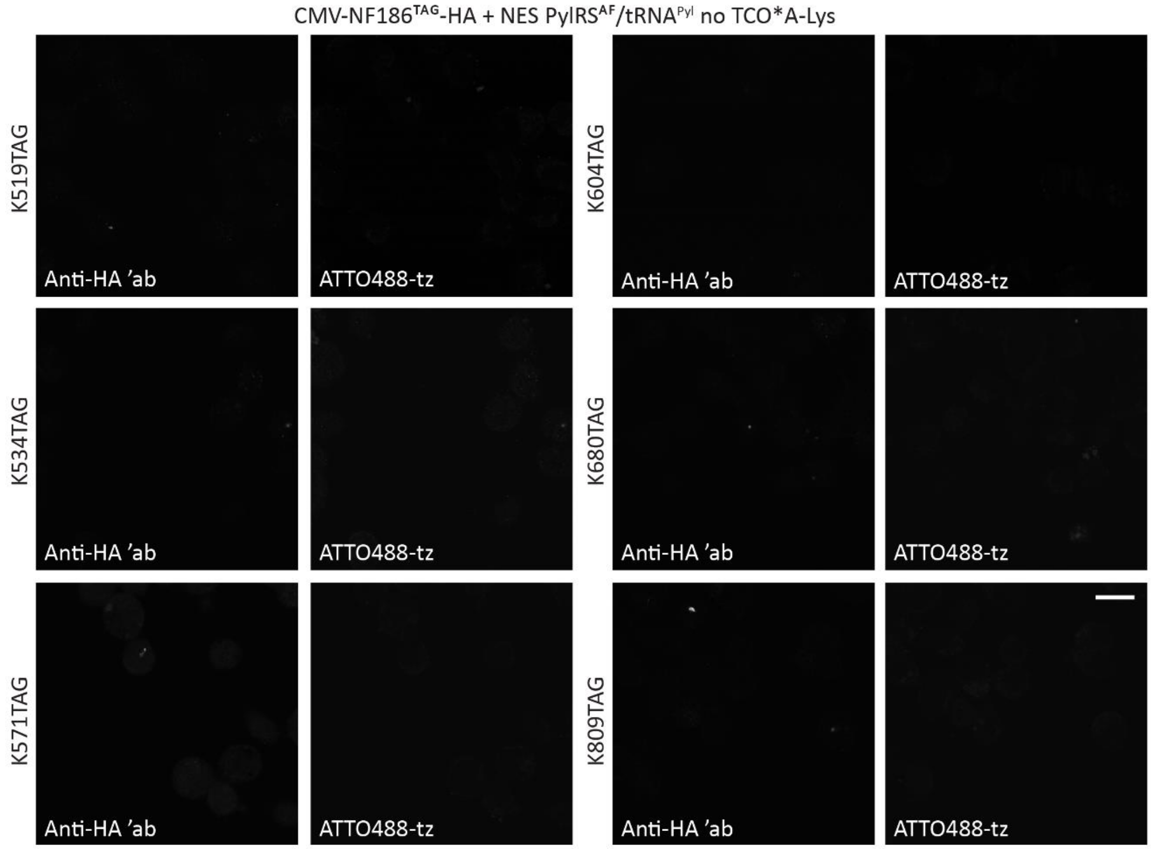


**Supplementary Fig. 2. Expression and click labeling of NF186$^{TAG}$-HA is not detected in the absence of the unnatural amino acid TCO*A-Lys in ND7/23 cells.** ND7/23 cells were co-transfected with NES PylRS$^{AF}$/tRNA$^{Pyl}$ and one of the CMV-NF186$^{TAG}$-HA amber mutants (K519TAG, K534TAG, K571TAG, K604TAG, K680TAG, and K809TAG) in the absence of the unnatural amino acid TCO*A-Lys. A day after the transfection, the cells were click-labeled with ATTO488 tetrazine (tz), fixed, and immunostained with anti-HA primary and Alexa Fluor Plus 647-conjugated secondary antibodies. Single-plane images were acquired with a confocal scanning microscope. Scale bar: 20 µm.

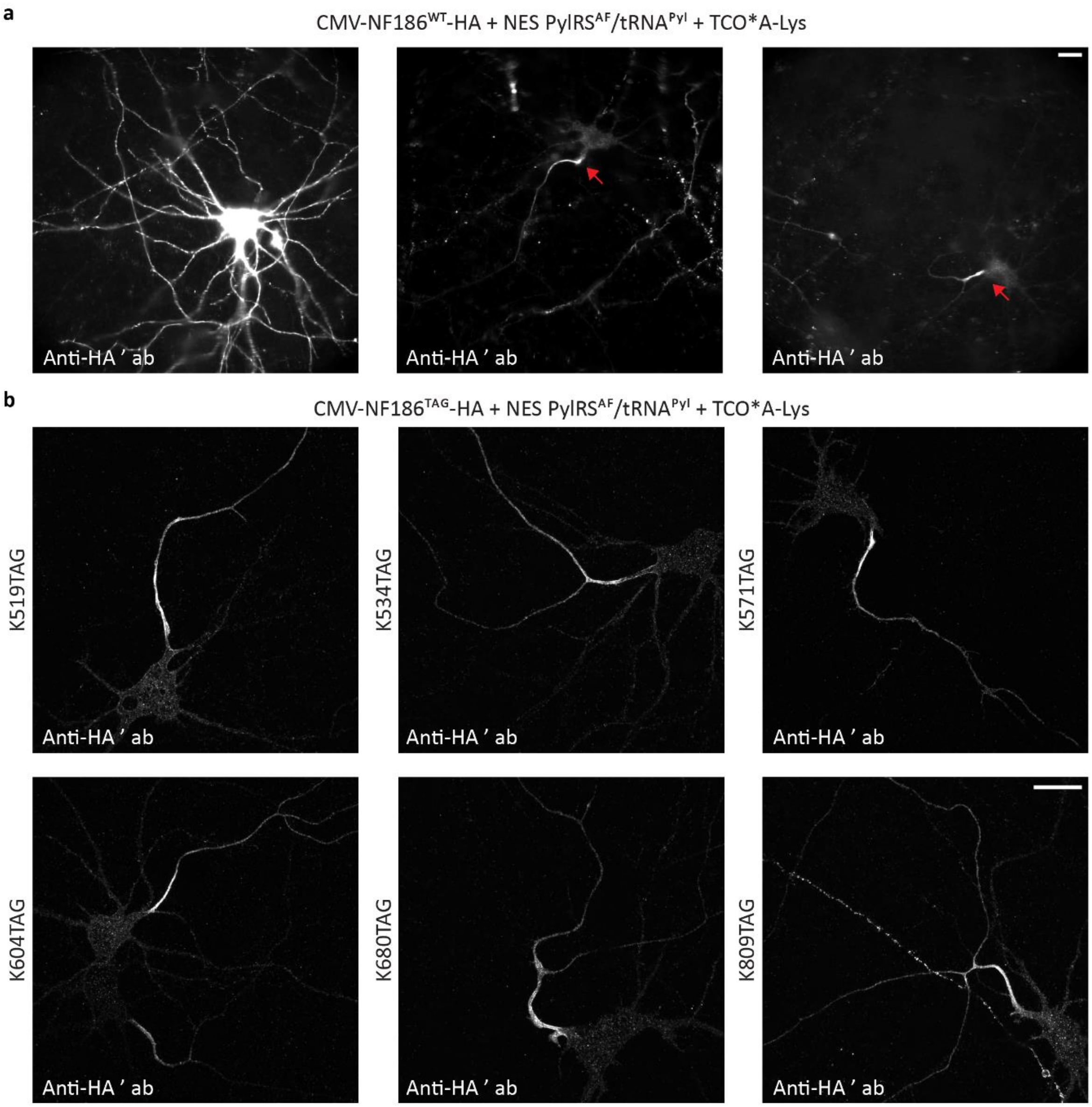


**Supplementary Fig. 3. Frequent mislocalization of CMV-NF186$^{WT}$-HA and CMV-NF186$^{TAG}$-HA amber mutants when expressed in primary neurons.** Representative images of primary rat cortical neurons at DIV 11 co-expressing NES PylRS$^{AF}$/tRNA$^{Pyl}$ and (**a**) CMV-NF186$^{WT}$-HA or (**b**) one of the CMV-NF186$^{TAG}$-HA amber mutants (K519TAG, K534TAG, K571TAG, K604TAG, K680TAG, and K809TAG) in the presence of the unnatural amino acid TCO*A-Lys. Four days after the transfection, the neurons were fixed and immunostained with anti-HA primary and Alexa Fluor 555-conjugated secondary antibodies.

**a**: Three different examples of neurons that expressed CMV-NF186$^{WT-}$HA. Most of the transfected neurons had an NF186 signal in all the processes (see the example image on the left panel). The arrows indicate axon initial segments.

**b**: In the neurons that expressed clickable CMV-NF186$^{TAG}$-HA, an NF186 signal was frequently observed along distal axons.

Neurons that expressed NF186$^{WT}$-HA were imaged with widefield microscopy (**a**), while amber mutants were imaged on a confocal scanning microscope (**b**). The Z-stack images are shown as maximum intensity projections. Scale bars: 20 µm (**a**–**b**).

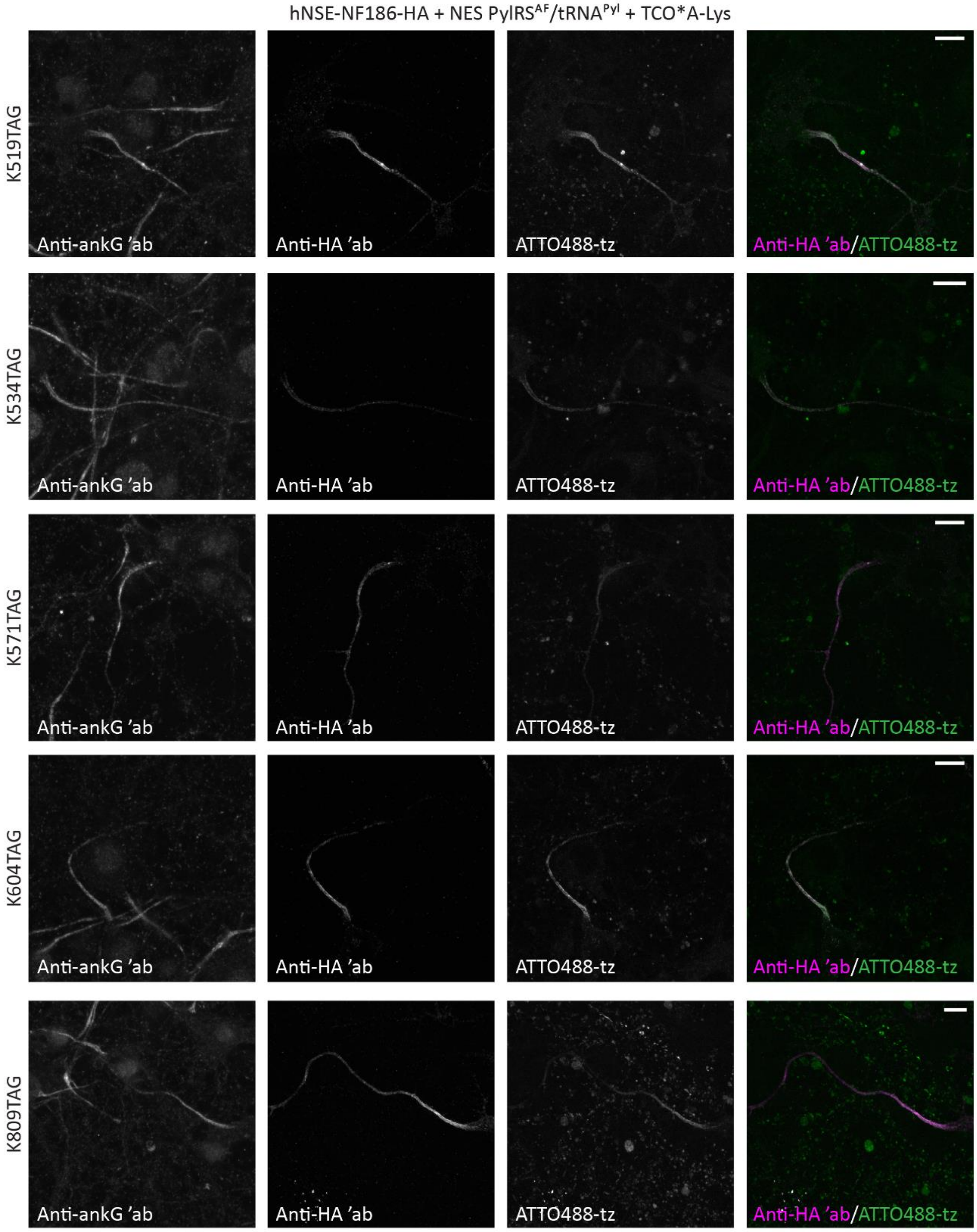


**Supplementary Fig. 4. Click labeling of additional hNSE-NF186$^{TAG}$-HA mutants in living primary neurons.** Representative images of primary rat cortical neurons at DIV 11 co-expressing NES PylRS$^{AF}$/tRNA$^{Pyl}$ and one of the hNSE-NF186$^{TAG}$-HA amber mutants (K519TAG, K534TAG, K571TAG, K604TAG, and K809TAG) in the presence of the unnatural amino acid TCO*A-Lys. The neurons were click-labeled with ATTO488-

tetrazine (tz), fixed and immunostained with anti-HA and anti-ankyrin G (ankG) primary and Alexa Fluor 555- and AF633-conjugated secondary antibodies. The Z-stack images are shown as maximum intensity projections. Scale bars: 10 µm.

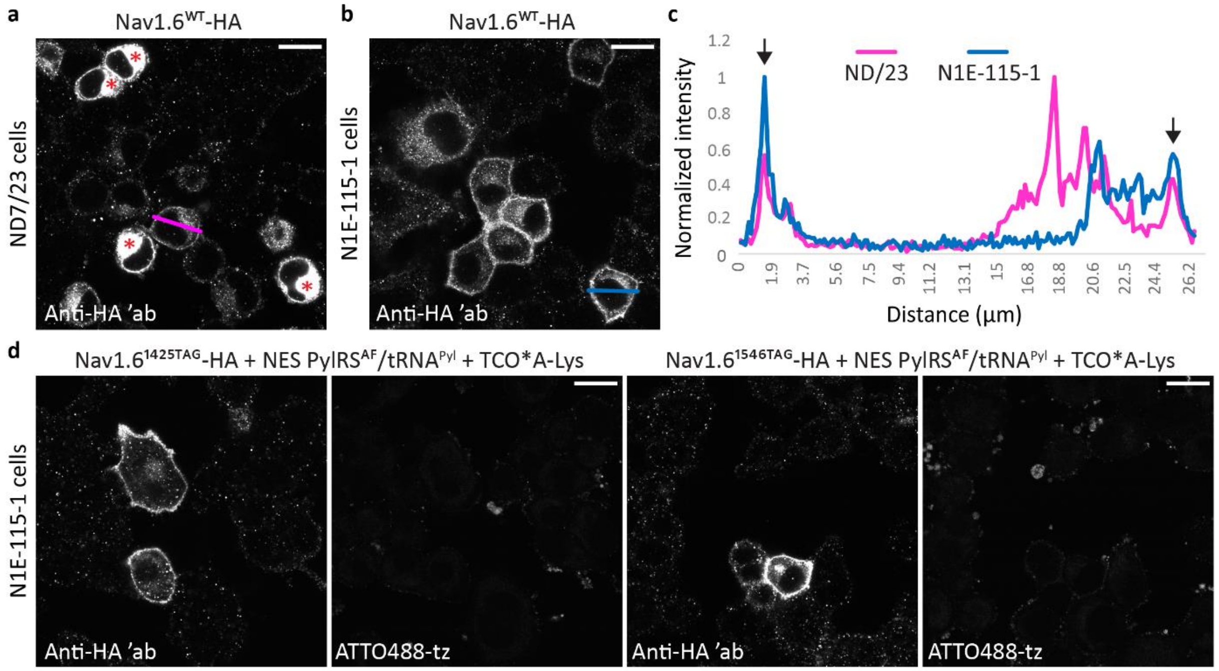


**Supplementary Fig. 5. Genetic code expansion and click labeling of $Na_V$1.6-HA in neuronal cell lines.**

**a**–**b**: Representative images of (**a**) ND7/23 and (**b**) N1E-115-1 cells co-expressing NES PylRS$^{AF}$/tRNA$^{Pyl}$, mouse β1- and β2 subunits, and $Na_V1.6^{WT}$-HA. Two days after the transfection, the cells were fixed and immunostained with anti-HA primary and Alexa Fluor (AF)555-conjugated secondary antibodies. The asterisks (in red) indicate ND7/23 cells with a strong HA signal in the cytoplasm and low signal on the cell membrane.

**c**: Graph of normalized line profile fluorescence intensity measurements for the HA signal on the membrane and in the cytoplasm of the ND7/23 or N1E-115-1 cells in panels a and b (pink and blue lines, respectively). The arrows in the graph indicate signals on the cell membrane.

**d**: N1E-115-1 cells co-expressing NES PylRS$^{AF}$/tRNA$^{Pyl}$, mouse β1- and β2 subunits, and $Na_v1.6^{TAG}$-HA in the presence of the unnatural amino acid TCO*A-Lys. Two days after the transfection, the cells were click-labeled with ATTO488-tz, fixed, and immunostained with anti-HA primary and AF555-conjugated secondary antibodies.

Single-plane images were acquired with a confocal scanning microscope. Scale bars: 20 µm (**a**, **b** and **d**).

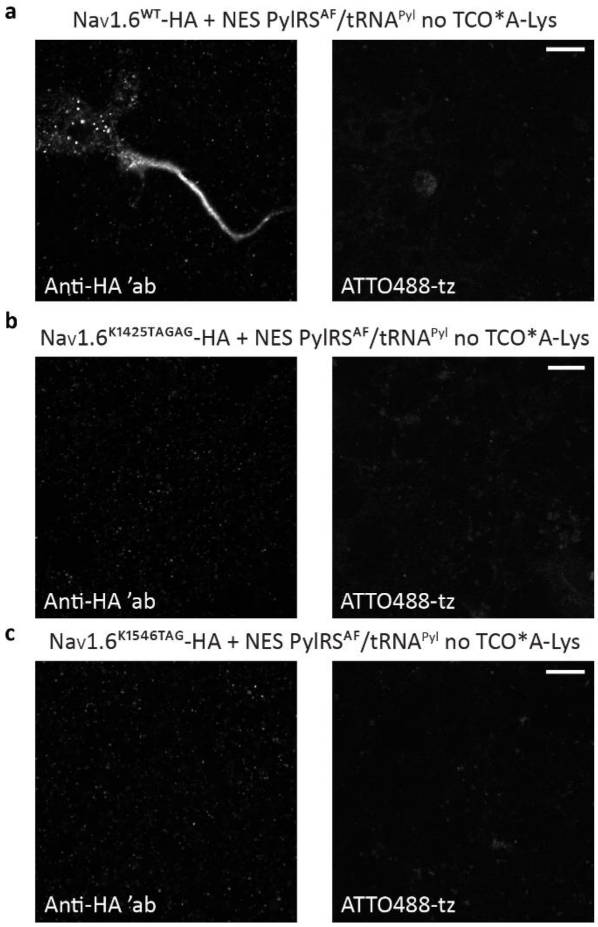


**Supplementary Fig. 6. Expression and click labeling of $Na_V$1.6$^{TAG}$-HA is not detected in primary neurons in the absence of the unnatural amino acid TCO*A-Lys.** Representative images of primary rat cortical neurons at DIV 12 co-expressing NES PylRS$^{AF}$/tRNA$^{Pyl}$ and (**a**) $Na_V$1.6$^{WT}$-HA, or (**b**) $Na_V$1.6$^{K1425TAG}$-HA (**c**) $Na_V$1.6$^{K1546TAG}$-HA in the absence of TCO*A-Lys. Neurons were click-labeled with ATTO488-tetrazine (tz), fixed, and immunostained with anti-HA primary and Alexa Fluor 555-conjugated secondary antibodies.

**a**: Neurons that expressed Nav1.6$^{WT}$-HA in the absence of TCO*A-Lys showed no click labeling with ATTO488-tz.

**b**–**c**: Expression and click labeling of $Na_V$1.6$^{K1425TAG}$-HA or $Na_V$1.6$^{K1546TAG}$-HA with ATTO488-tz were not detected in the absence of TCO*A-Lys.

The Z-stack confocal images (**b**, **c**) are shown as maximum intensity projections. Scale bars: 10 µm (**a**–**c**).

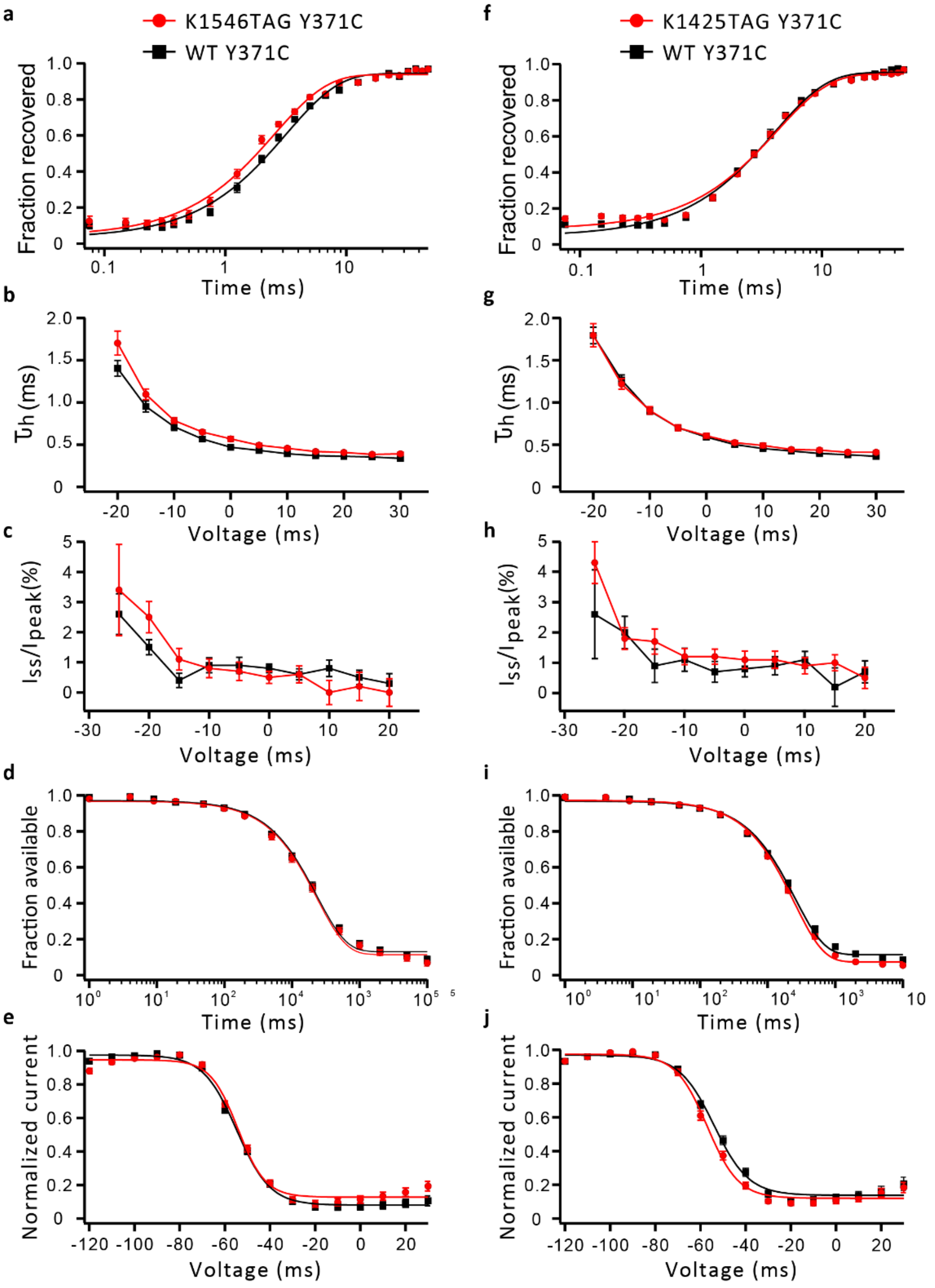


**Supplementary Fig. 7. Additional biophysical properties of Na$_V$1.6$^{WT/TAG}$-HA in N1E-115-1 cells.** (**a**–**e**) N1E-115-1 cells were co-transfected with NES PylRS$^{AF}$/tRNA$^{Pyl}$**,** multigene plasmid encoding mouse β1- and β2 subunits and GFP, and Na$_V$1.6$^{WT,\ Y371C}$-HA or Na$_V$1.6$^{K1546TAG,\ Y371C}$-HA. (**f**–**j**) N1E-115-1$^{β1β2}$ stable cells were co-transfected with NES PylRS$^{AF}$/tRNA$^{Pyl}$, Na$_V$1.6$^{WT,\ Y371C}$-P2A-eGFP, or Na$_V$1.6$^{K1425TAG,\ Y371C}$-P2A-eGFP. Na$^+$

currents were recorded two days after the transfection in the presence of 500 nM tetrodotoxin (TTX) that blocked endogenous $Na^+$ currents.

**a, f:** Time course of recovery from fast inactivation at -100 mV. K1546TAG significantly accelerated the recovery from fast inactivation compared to the WT channels.

**b, g:** Voltage dependence of the major time constant of fast inactivation $\tau_h$. K1546TAG significantly delayed the transition from activation to fast inactivation compared to the WT channels.

**c, h:** Voltage dependence of the persistent current ($I_{ss}/I_{peak}$).

**d, i:** Entry into slow inactivation. The lines represent fits of a first-order exponential function to the data points.

**e, j:** Steady-state slow inactivation. The lines represent Boltzmann functions fit to the data points. Shown are means ± standard errors of the mean (SEMs).

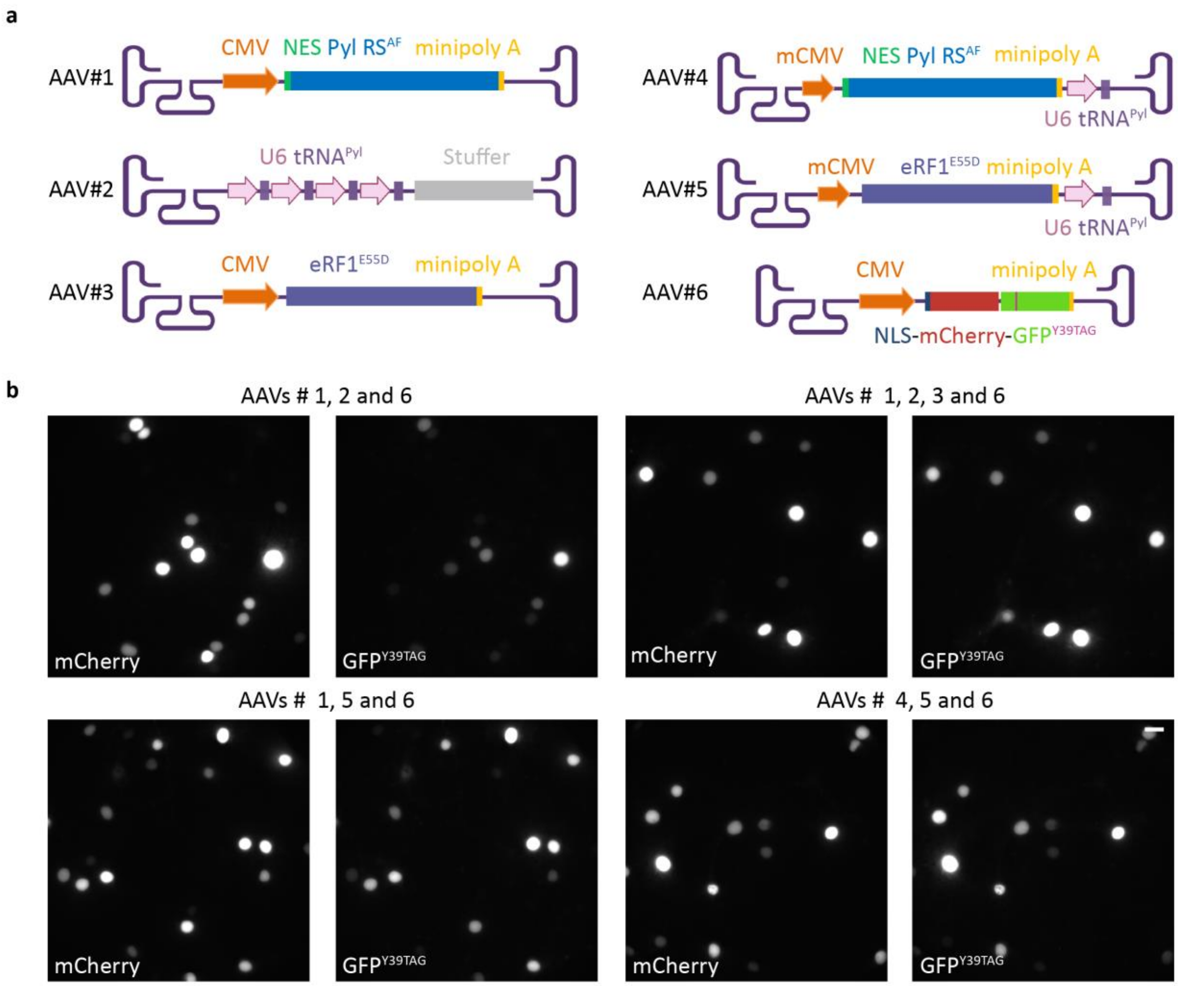


**Supplementary Fig. 8. Adeno-associated virus (AAV)-based vectors enabling efficient genetic code expansion of a fluorescent reporter in primary neurons.**

**a**: **S**chematic representation of AAV-based viral vectors used for transduction of primary rat cortical neurons. In addition to AAV9A2 carrying a (NLS)-mCherry-GFP$^{Y39TAG}$ fluorescent reporter cassette (AAV#6), we developed AAV9A2 vectors with various combinations of genes for genetic code expansion: AAV#1: NES PylRS$^{AF}$ expressed from a CMV promoter; AAV#2: four copies of tRNA$^{Pyl}$ expressed from a U6 promoter; AAV#3: eukaryotic release factor eRF1$^{E55D}$ expressed from a CMV promoter; AAV#4: NES PylRS$^{AF}$ expressed from a minimal (min)CMV promoter and one copy of tRNA$^{Pyl}$ expressed from a U6 promoter; and AAV#5: minCMV eRF1$^{E55D}$ and one copy of U6 tRNA$^{Pyl}$.

**b**: Representative widefield images of primary neurons at DIV 11 co-expressing NLS-mCherry-GFP$^{Y39TAG}$ (AAV#6) and different combinations of AAVs (AAV#1-5) for genetic code expansion in the presence of the unnatural amino acid TCOA*-Lys. Scale bars: 20µm. The schemes in panel a were made in BioRender.com.

#### Supplementary Tables

**Supplementary Table 1.** Pairwise comparisons of the AIS length (measured in ankG channels) between NF186 transfected (HA+) and surrounding untransfected (HA-) neurons using the independent-samples Kruskal-Wallis test.

Groups: 1. WT HA+; 2. WT HA-; 3. K519TAG HA+; 4. K519TAG HA-; 5. K604TAG HA+; 6. K604TAG HA-; 7. K680TAG HA+; and 8. K680TAG HA-. Pairwise comparisons of transfected and untransfected neurons for each of the constructs (1-2, 3-4, 5-6, and 7-8) are highlighted in grey.

| Pairwise comparisons | | | | | |
|---|---|---|---|---|---|
| Comparisons | Test statistic | Standard error | Standard test statistic | Significance | Adjusted significance [a] |
| 4.00-2.00 | 6.433 | 11.390 | 0.565 | 0.572 | 1.000 |
| 4.00-8.00 | -9.382 | 12.969 | -0.723 | 0.469 | 1.000 |
| 4.00-6.00 | -23.138 | 17.448 | -1.326 | 0.185 | 1.000 |
| 4.00-1.00 | 40.443 | 13.875 | 2.915 | 0.004 | 0.100 |
| 4.00-7.00 | -41.832 | 13.875 | -3.015 | 0.003 | 0.072 |
| 4.00-3.00 | 42.133 | 16.240 | 2.594 | 0.009 | 0.265 |
| 4.00-5.00 | -52.360 | 16.795 | -3.118 | 0.002 | 0.051 |
| 2.00-8.00 | -2.949 | 11.693 | -0.252 | 0.801 | 1.000 |
| 2.00-6.00 | -16.705 | 16.523 | -1.011 | 0.312 | 1.000 |
| 2.00-1.00 | 34.010 | 12.691 | 2.680 | 0.007 | 0.206 |
| 2.00-7.00 | -35.399 | 12.691 | -2.789 | 0.005 | 0.148 |
| 2.00-3.00 | -35.700 | 15.241 | -2.342 | 0.019 | 0.537 |
| 2.00-5.00 | -45.927 | 15.831 | -2.901 | 0.004 | 0.104 |
| 8.00-6.00 | 13.756 | 17.648 | 0.779 | 0.436 | 1.000 |
| 8.00-1.00 | 31.062 | 14.125 | 2.199 | 0.028 | 0.781 |
| 8.00-7.00 | 32.450 | 14.125 | 2.297 | 0.022 | 0.605 |
| 8.00-3.00 | 32.751 | 16.455 | 1.990 | 0.047 | 1.000 |
| 8.00-5.00 | 42.978 | 17.002 | 2.528 | 0.011 | 0.321 |
| 6.00-1.00 | 17.306 | 18.325 | 0.944 | 0.345 | 1.000 |
| 6.00-7.00 | -18.694 | 18.325 | -1.020 | 0.308 | 1.000 |
| 6.00-3.00 | 18.995 | 20.175 | 0.942 | 0.346 | 1.000 |
| 6.00-5.00 | 29.222 | 20.624 | 1.417 | 0.157 | 1.000 |
| 1.00-7.00 | -1.389 | 14.962 | -0.093 | 0.926 | 1.000 |
| 1.00-3.00 | -1.689 | 17.178 | -0.098 | 0.922 | 1.000 |
| 1.00-5.00 | -11.917 | 17.703 | -0.673 | 0.501 | 1.000 |

| | | | | | |
|---|---|---|---|---|---|
| 7.00-3.00 | 0.301 | 17.178 | 0.017 | 0.986 | 1.000 |
| 7.00-5.00 | 10.528 | 17.703 | 0.595 | 0.552 | 1.000 |
| 3.00-5.00 | -10.227 | 19.612 | -0.521 | 0.602 | 1.000 |

*Note.* Each row tests the null hypothesis that the Sample 1 and Sample 2 distributions are the same. Asymptotic significances (two-sided tests) are displayed. The significance level is 0.05. [a.] Significance values have been adjusted by the Bonferroni correction for multiple tests.

**Supplementary Table 2.** Multiple comparisons of the AIS length (measured in ankG channels) between $Na_V1.6^{TAG}$-HA transfected (HA+) and surrounding untransfected (HA-) cells using one-way ANOVA with the Tukey *posthoc*.

Groups: 1. K1425TAG HA+; 2. K1425TAG HA-; 3. K1546TAG HA+; and 4. K1546TAG HA-. Multiple comparisons of transfected and untransfected neurons for each of the constructs (1-2, 3-4) are highlighted in grey.

| **Multiple comparisons** | | | | | | |
|---|---|---|---|---|---|---|
| Tukey HSD | | | | | | |
| (I) Category | (J) Category | Mean difference (I-J) | Stdandard error | Significance | 95% Confidence interval | |
| | | | | | Lower bound | Upper bound |
| 1.00 | 2.00 | 3.66061 | 1.64177 | 0.124 | -0.6418 | 7.9630 |
| | 3.00 | 0.78047 | 1.96907 | 0.979 | -4.3797 | 5.9406 |
| | 4.00 | 4.90039* | 1.79007 | 0.037 | 0.2094 | 9.5914 |
| 2.00 | 1.00 | -3.66061 | 1.64177 | 0.124 | -7.9630 | 0.6418 |
| | 3.00 | -2.88014 | 1.74582 | 0.357 | -7.4552 | 1.6949 |
| | 4.00 | 1.23979 | 1.54110 | 0.852 | -2.7988 | 5.2784 |
| 3.00 | 1.00 | -0.78047 | 1.96907 | 0.979 | -5.9406 | 4.3797 |
| | 2.00 | 2.88014 | 1.74582 | 0.357 | -1.6949 | 7.4552 |
| | 4.00 | 4.11993 | 1.88595 | 0.136 | -0.8224 | 9.0622 |
| 4.00 | 1.00 | -4.90039* | 1.79007 | 0.037 | -9.5914 | -0.2094 |
| | 2.00 | -1.23979 | 1.54110 | 0.852 | -5.2784 | 2.7988 |
| | 3.00 | -4.11993 | 1.88595 | 0.136 | -9.0622 | 0.8224 |

*Note.* *. The mean difference is significant at the 0.05 level.

**Supplementary Table 3.** Biophysical properties of $Na_V1.6^{WT,\ Y371C}$-HA and $Na_V1.6^{K1546TAG,\ Y371C}$-HA recorded in neuronal N1E-115-1 cells, or $Na_V1.6^{WT,\ Y371C}$-P2A-eGFP and $Na_V1.6^{K1425TAG,\ Y371C}$-P2A-eGFP recorded in neuronal N1E-115-1$^{\beta1\beta2}$ stable cells.

| | **Steady- state activation** | | | **Steady- state inactivation** | | | **$\tau_h$ at 0 mV (ms)** | **n** | **$\tau_{rec}$ at -100 mV (ms)** | **n** | **Current density (pA/pF)** | **n** |
|---|---|---|---|---|---|---|---|---|---|---|---|---|
| | **$V_{1/2}$ (mV)** | **k** | **n** | **$V_{1/2}$ (mV)** | **k** | **n** | | | | | | |
| **$Na_V1.6^{WT}$** | -9.9±1.1 | -7.3±0.3 | 18 | -58.8±0.5 | 4.9±0.1 | 18 | 0.59±0.02 | 18 | 3.6±0.2 | 17 | -124.6±15.5 | 18 |
| **$Na_V1.6^{K1425TAG}$** | -11.4±0.6 | -7.4±0.2 | 20 | -60.2±0.5 | 4.8±0.1 | 20 | 0.60±0.03 | 20 | 3.6±0.2 | 20 | -89.0±5.5* (p=0.0426) | 20 |
| **$Na_V1.6^{WT}$** | -11.8±1.0 | -6.8±0.3 | 20 | -58.6±0.5 | 4.8±0.1 | 20 | 0.47±0.02 | 20 | 2.8±0.1 | 20 | -149.4±17.5 | 20 |
| **$Na_V1.6^{K1546TAG}$** | -11.5±0.8 | -6.6±0.3 | 18 | -55.8±0.6** (p= 0.0025) | 4.9±0.1 | 17 | 0.57±0.03** (p=0.0026) | 18 | 2.4±0.2* (p=0.0173) | 18 | -105.5±12.5 | 18 |

*Note.* Data are presented as means ± standard errors of the mean (SEMs); number of recorded cells (n); * $p < 0.05$, ** $p < 0.01$, Student´s *t*-test or Mann-Whitney U test.

**Supplementary table 4.** Mean AIS fluorescence intensity (measured in click channels) between control ($Na_V1.6^{K1546TAG,\ Y371C}$-HA) and loss-of-function variants ($Na_V1.6^{K1546TAG,\ Y371C.I1652N}$-HA and $Na_V1.6^{K1546TAG,\ Y371C,\ T1785P}$-HA) using the independent-samples Kruskal-Wallis test.

| **Independent-samples Kruskal-Wallis test: summary** | |
|---|---|
| Total number of cells (n) | 56 |
| Test statistic | 3.135[a,b] |
| Degree of freedom | 2 |
| Asymptotic significance (2-sided test) | 0.209 |

[a] The test statistic is adjusted for ties. [b] Multiple comparisons are not performed because the overall test does not show significant differences across samples.

**Supplementary Table 5.** Mutagenesis and cloning primers

| Purpose | Primer name | Primer sequence 5'→3' |
|---|---|---|
| Deletion of HA tag from the N terminus of NF186 | HA(del)-NF186fw | GCCATTGAGATTCCGATGGATCCAAGCATTCAGAATGAG |
| | HA(del)-NF186rv | CTCATTCTGAATGCTTGGATCCATCGGAATCTCAATGGC |
| NF186$^{TAG}$ mutagenesis | NF186$^{K519TAG}$fw | GAGGTCTAGGACCCCACCAGGATCTACAGGATG |
| | NF186$^{K519TAG}$rv | GGGGTCCTAGACCTCCAGGCGGACTTGATTTTCA |
| | NF186$^{K534TAG}$fw | GTGGCCTAGAGGGGCACCACAGTGCAG |
| | NF186$^{K534TAG}$rv | GCCCCTCTAGGCCACCTGGTCTTCAGGC |
| | NF186$^{K571TAG}$fw | AGGATGTAGAAGGAAGATGACTCCCTGACCATCTTCG |
| | NF186$^{K571TAG}$rv | TTCCTTCTACATCCTGTTTCCAATGTAGAGTGGCTC |
| | NF186$^{K604TAG}$fw | CTGGCATAGGCCTACCTCACTGTTCTAGCTGATCAG |
| | NF186$^{K604TAG}$rv | GTAGGCCTATGCCAGGTCCTGGTCCAG |
| | NF186$^{K680TAG}$fw | CACTCCTAGTTCCCAGGCAGTGTCAACTCAG |
| | NF186$^{K680TAG}$rv | TGGGAACTAGGAGTGGTCATGCCAGACTCCT |
| | NF186$^{K809TAG}$fw | TTTGGGTAGGGCCCGGAGCCTGAAAC |
| | NF186$^{K809TAG}$rv | CGGGCCCTACCCAAAGTCATTTTCAGCCTGGACTC |
| Addition of HA tag to the C terminus of NF186 | NF186-HAfw | GGTGGTGGGCCCTGAAGACCCCAAAGAAG |
| | NF186-HArv | ACCACCGCGGCCGCTCAAGCGTAGTCTGGGACGTCGTATG GGTAGGCCAGGGAATAGATGGCATTGACTG |
| Cloning of hNSE into NF186 plasmid | hNSE(AseI)fw | GGTGGTATTAATTGTATGCAGCTGGACCTAGGAGAG AAG |
| | hNSE(BglII)rv | ACCACCAGATCTCGGTGGTAGTGGCGG |
| Cloning of *mSCN1B* into pACEMam2 to make multigene plasmid | *mSCN1B*(KpnI) rv | GGTGGTGCTAGCCACCATGGGGACGCTGCTGGCTCT |
| | *mSCN1B*(NheI)fw | ACCACCGGTACCTTATTCAGCCACCTGGACGCCT |
| Cloning of *mSCN2B* into pMDS to make multigene plasmid | *mSCN2B*(NheI)fw | GGTGGTGCTAGCCACCATGGAAGTCACAGCGCCCACC A |
| | mSCN2B(KpnI)rv | ACCACCGGTACCTTACTTGGTGCCATCTTCCGCGTTG |
| Cloning of mGFP into pMDC to make multigene plasmid | mGFP(BamHI)fw | GGTGGTGGATCCCACCATGGTGAGCAAGGGCG |
| | mGFP(XbaII)rv | ACCACCTCTAGATTACTTGTACAGCTCGTCCATGCCGA |
| Cloning of *mSCN1B* and *mSCN2B* for generation of N1E-115-1$^{β1β2}$ | *mSCN1B*(NheI)fw | GGTGGTGCTAGCCACCATGGGGACGCTGCTGGCTCT |
| | *mSCN1B*(NotI)rv | CCGTCCGCGGCCGCCTATTCAGCCACCTGGACGCC |
| | *mSCN2B*(NheI)fw | GGTGGTGCTAGCCACCATGCACAGGGATGCCTGGCTACC |
| | *mSCN2B*(NotI)rv | CCGTCCGCGGCCGCTTACTTGGTGCCATCTTCCGCGT |
| Amplification of *mSCN1B* and *mSCN2B* from N1E-1151$^{β1β2}$ gDNA | *mSCN1B*rv | CTATTCAGCCACCTGGACGCCTG |
| | *mSCN2B*rv | TTACTTGGTGCCATCTTCCGCGTTG |
| | PiggyBACfw | ATGTAATTACGTCCCTCCCCCGCTAG |
| Loss-of-function *SCN8A* mutagenesis | *mSCN8A*$^{I652N}$rv | CAGCAGGCCGTTATTGAACAGGGCTGGCAG |
| | *mSCN8A*$^{I652N}$fw | CTGCCAGCCCTGTTCAATAACGGCCTGCTG |
| | *mSCN8A*$^{T1785P}$rv | CCAGATCTCATAGAAGGGCTCGAAGTCATCCTCAG |
| | *mSCN8A*$^{T1785P}$fw | CTGAGGATGACTTCGAGCCCTTCTATGAGATCTGG |

| | | |
|---|---|---|
| Introduction of TTXr mutation into *mSCN8A* | *mSCN8A*$^{Y371C}$fw | GTACAGGTTCTCCCAGCAGTCCTGGGTCATCAG |
| | *mSCN8A*$^{Y371C}$rv | CTGATGACCCAGGACTGCTGGGAGAACCTGTAC |
| Addition of HA tag on the C terminus of $Na_V1.6$ | *mSCN8A*-HAfw | (Phos)CTAGATACCCATACGACGTCCCAGACTACGCTTAAG GGCC |
| | *mSCN8A*-HArv | (Phos)CTTAAGCGTAGTCTGGGACGTCGTATGGGTAT |
| Measurement of the AAV titer via ddPCR | GFPfw | ATCTTCTTCAAGGACGACG |
| | GFPrv | TCCTCCTTGAAGTCGATGC |
| | Probe | FAM-ACGACGGCAACTACA-BHQ1 |
